# Developmental diet alters the fecundity-longevity relationship and age-related gene expression in *Drosophila melanogaster*

**DOI:** 10.1101/2023.01.16.524185

**Authors:** David H. Collins, David C. Prince, Jenny L. Donelan, Tracey Chapman, Andrew F. G. Bourke

## Abstract

The standard evolutionary theory of ageing predicts a negative relationship (trade-off) between fecundity and longevity. However, this relationship can become positive: (i) under the influence of longevity-enhancing mutations; (ii) when individuals have unequal resources; and (iii) in eusocial insects, in which reproductive queens outlive less- or non-reproductive workers. Developmental diet is likely to be central to determining trade-offs as it affects key fitness traits such as adult body size, but its exact role remains uncertain. For example, in *Drosophila melanogaster* fruit flies, changes in adult diet can affect fecundity, longevity, and gene expression throughout life, but it is unknown how changes in developmental (larval) diet affect fecundity-longevity relationships or gene expression in adults. Using *D. melanogaster*, we therefore tested the hypothesis that variation in developmental diet alters the directionality of fecundity-longevity relationships in adults, and characterised associated gene expression changes. We reared *D. melanogaster* larvae on low (20%), medium (100%), and high (120%) SYA (Sugar Yeast Agar) diets, and transferred adult females developing from these larvae to a common (110% SYA) adult diet. We measured life-time fertility (realised fecundity) and longevity of individual adult females and, using mRNA-seq, profiled gene expression changes across two time-points. Adult females raised on the different larval diets exhibited fecundity-longevity relationships that were significantly different in directionality, i.e., varied from positive to negative, despite minimal differences in mean life-time fertility or longevity. Treatments also differed in age-related gene expression, including expression of genes known to be associated with ageing. Hence, this study shows that the sign of fecundity-longevity relationships in adult insects can be altered and even reversed by variation in larval diet quality. Furthermore, larval diet quality may be a key mechanistic factor underpinning positive fecundity-longevity relationships observed in species such as eusocial insects.

## Introduction

The standard evolutionary theory of ageing predicts that as individuals grow older, the strength of selection for increased survivorship declines with age^1^. Therefore, individuals suffer the age-related decrease in performance and survivorship that defines ageing (senescence)^2^. At the same time, because individuals have finite resources available to them, they should optimise relative investment between reproduction and somatic maintenance to maximise fitness^3^. This causes trade-offs between reproduction and longevity^4,5^ with elevated reproduction often incurring costs to longevity (the costs of reproduction)^6^. Such trade-offs and costs are evident in the negative relationship between fecundity and longevity observed in most species.

Although a negative fecundity-longevity relationship is typical, fecundity and longevity can become uncoupled and their relationship can be reversed^7^. That is, some species or populations may exhibit positive fecundity-longevity relationships^4^, and this can occur for several reasons. First, in *Drosophila melanogaster* mutations can increase longevity without apparent reproductive costs^8–12^, particularly for mutations in the highly evolutionarily conserved insulin/insulin-like growth factor signalling (IIS) and target of rapamycin (TOR) network (IIS-TOR). This network regulates nutrient sensitivity and is an important component of ageing in several species, including nematodes, insects, and humans^2,13^.

Second, fecundity and longevity can become uncoupled when resource acquisition varies between individuals^14,15^. Within a population, this could translate into some particularly well-resourced individuals having higher fecundity and longevity than poorly-resourced individuals, giving the appearance of an uncoupling (or reversal) of the usual negative fecundity-longevity relationship. However, since costs of reproduction are not abolished even in well-resourced individuals^14,15^, a trade-off between fecundity and reproduction remains at the individual level.

Third, fecundity and longevity can become uncoupled within and between the castes of eusocial insects^16–23^, i.e., species with a long-lived caste specialising in reproduction (queens or kings) and a short-lived caste specialising in colony maintenance (workers), which is therefore less fecund or even sterile^24–26^. In some eusocial insects, reproductive individuals appear to have escaped the costs of reproduction completely^27–30^. As with long-lived *D. melanogaster* mutants, these individuals may have achieved a decoupling of fecundity and longevity through rewiring of the IIS-TOR network^13,31^, which forms part of the TI-J-LiFe network hypothesised to underpin ageing and longevity in eusocial insects by Korb et al.^32^. By contrast, the apparent lack of costs of reproduction may have been caused by asymmetric resourcing in some eusocial insects. That is, long-lived and highly reproductive queens and workers may be able to overcome the costs of reproduction by having access to higher quality resources as larvae^22,23^.

*D. melanogaster* is an important model species for exploring the effect of diet on longevity and fecundity. Dietary restriction in this species causes life-span extension in adults^33–36^ and larvae^37–40^. Differences in diet quality can also decouple fecundity-longevity relationships in *D. melanogaster* adults^36,41^. For example, longevity and reproduction are each maximised on different adult diets, with longevity maximised on a low protein:carbohydrate diet, and reproduction maximised on a high protein:carbohydrate diet^36,41^. Changes in adult diets are also accompanied by age-related transcriptional changes^41–43^. As is found for the long-lived mutant *D. melanogaster* strains, dietary effects might be underpinned by downregulation of the IIS-TOR signalling network^44^ While changes in developmental (larval) diet generally have less strong effects than changes in the adult diet, *D. melanogaster* larvae reared on restricted (but not starvation) diets exhibit longer development durations, lower development success, smaller body sizes, increased longevity and, in some situations, increased fecundity^38,39,45^. However, other studies have shown that low-quality larval diets reduced lifespan and/or fecundity^45,46^. Changes in larval diet and rearing environment can also interact with the adult diet to have contrasting effects on adult life-history traits^38,45^. Irrespective of whether a low-quality larval diet affects longevity and fecundity in a positive or a negative way, the outcomes of these studies predict that the fecundity-longevity relationship in adult flies could itself be affected by the quality of the larval diet. Specifically, variation in the larval diet is expected to be important in shaping the relationship between adult longevity and fecundity as it determines key fitness traits such as body size^45^.

However, to date no study has explored whether *D. melanogaster* that experience different larval diets show differences in the directionality of their fecundity-longevity relationship as adults. In particular, it is not known whether, when larval diet is manipulated, the negative fecundity-longevity relationship usually found in *D. melanogaster* females^4^ can be reversed so that it becomes positive. Establishing this is important because of the need to understand the mechanisms underpinning fecundity-longevity relationships and because larval diet could be a key (but untested) determinant of the directionality of the fecundity-longevity relationship in other species such as eusocial insects. Under this hypothesis, differences in the quality of the larval diet lead to populations of individuals varying in resource level, leading in turn to changes in the directionality of the fecundity-longevity relationship. For example, abundant larval resources may amplify genetic, environmental, or stochastic differences in individual quality. This could then cause high-quality individuals (which can capitalise on increased resource abundance as larvae) to exhibit high adult fecundity and longevity, and low-quality individuals (which cannot capitalise on increased resource abundance) to exhibit low fecundity and longevity, causing a positive fecundity-longevity relationship on high quality diets overall^22^. These trends are observed in eusocial insect queens, which receive superior larval diets compared to workers (so rendering them high-quality individuals) and exhibit a positive fecundity-longevity relationship^22^. Therefore, the hypothesis predicts that, other things equal, a superior larval diet causes insects to exhibit a positive fecundity-longevity relationship.

Accordingly, the first aim of this study was to determine the effect of larval diet on the directionality of the fecundity-longevity relationship in adult *D. melanogaster* females. To address this, we reared *D. melanogaster* larvae on three treatment diets: low-quality (20% SYA, i.e., Sugar Yeast Agar), medium-quality (100% SYA), and high-quality (120% SYA) diets (henceforth, L, M and H treatments, respectively). We then reared the adult female flies eclosing from all three treatment diets on a different, standardised, common garden adult diet (110% SYA) and, over the adult flies’ lifetimes, measured the effect of treatment (i.e. larval diet) on adult longevity (number of days lived post-eclosion) and fertility (i.e., realised fecundity, as measured by egg production and offspring production over each female’s lifetime). As larval diet affects development duration, development success, and adult body size, we also measured these traits. We conducted the experiment with females to allow direct measures of fertility, and because empirical and theoretical evidence shows that the lifehistory traits of females are more responsive to diet than those of males^45^. We hypothesised that, in addition to affecting adult female longevity and fertility (as other studies have shown), the differences in larval diet would change the directionality of the fecundity-longevity relationship within each treatment. More specifically, we hypothesised that the fecundity-longevity relationship would be either positive, or less negative, in the H females (reared on a high-quality larval diet) compared to the M and L females (reared on medium and low-quality larval diets, respectively).

The second, related aim of this study was to determine whether a change in the directionality of the fecundity-longevity relationship would have an effect on age-related gene expression in *D. melanogaster* females. *D. melanogaster* females are known to show age-related transcriptional changes that covary with reproductive senescence^42,43^ and adult diet^41^. However, the genetic correlates of changes in the directionality of the fecundity-longevity relationship following manipulation of larval diet have not previously been investigated. We therefore used mRNA-seq to sequence transcriptomes from head, fat body (abdomen minus gut and ovaries), and ovaries of females from the M and H treatments (as these two treatments showed the most pronounced difference in the directionality of the fecundity-longevity relationship) at two relative time intervals, i.e., after 10% and 60% of females in each treatment had died. We hypothesised that a change in the directionality of the fecundity-longevity relationship would alter the pattern of gene-expression change with relative age in each of these treatments. As the IIS-TOR network has been hypothesised to be influential in underpinning the changes in fecundity-longevity relationships seen in long-lived *D. melanogaster* mutants and in queens of eusocial insects, we also tested whether components of this network were differentially expressed with age in *D. melanogaster* depending on the directionality of the fecundity-longevity relationship.

## Methods

### Fly rearing and sample collection

The overall design of the experiment was to rear females on different larval diets, maintain them on a common adult diet, mate them repeatedly (once every eight days) until death to allow them to remain in egg-laying condition, and then to test the effect of larval diet on individual adult female longevity, fertility, and age-related gene expression (Figure 1).

**Figure 1.**
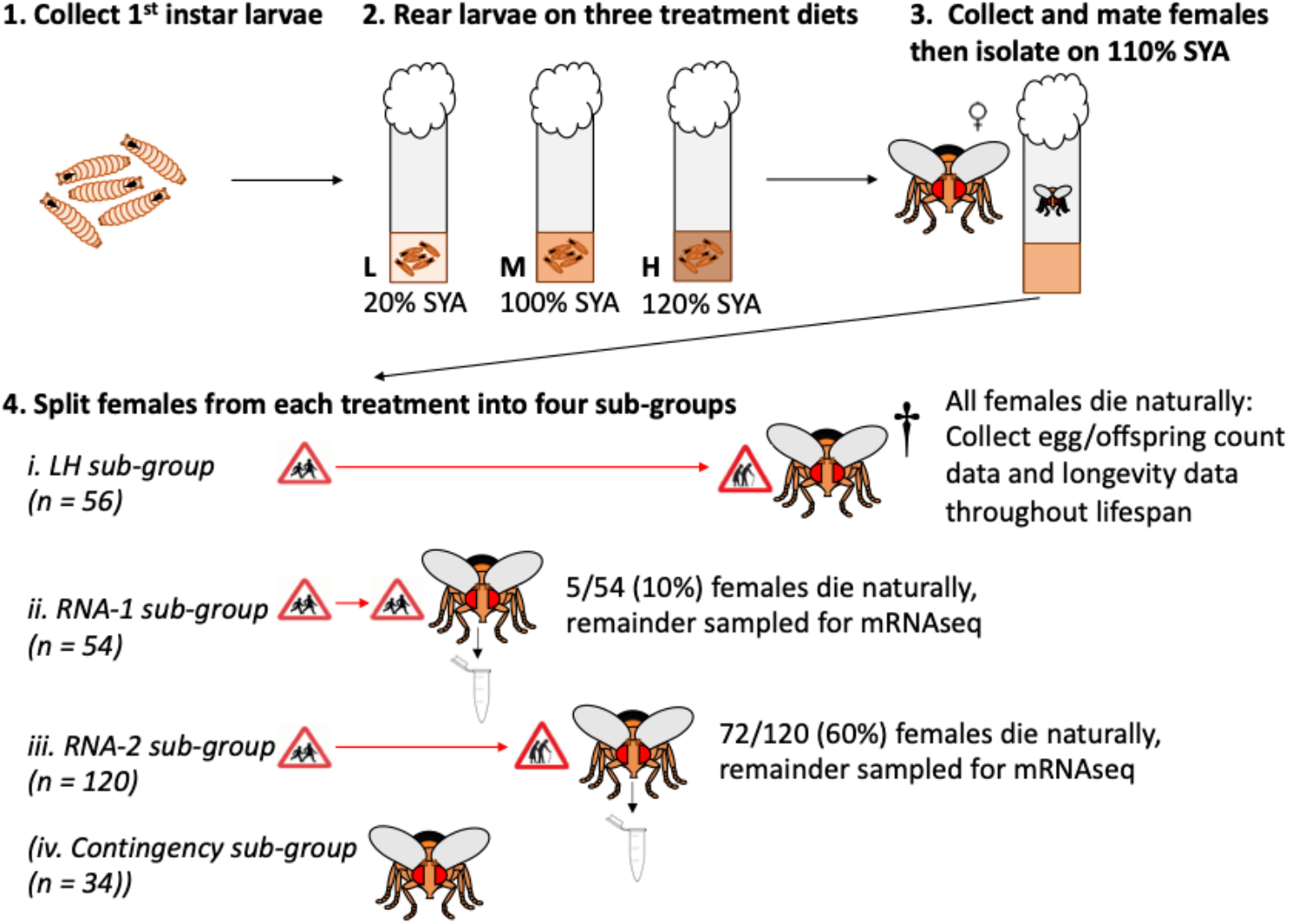
Outline of experimental design to determine the effect of larval diet (treatment) on the fecundity-longevity relationship and age-related gene expression of adult female *Drosophila melanogaster*. Flies were reared as larvae (of both sexes) on three treatment diets: L: low quality, 20% SYA; M: medium quality, 100% SYA; H: high quality, 120% SYA. Adult males were discarded or used for mating and adult females, after mating, were maintained on a common 110% SYA diet. Females were split into four sub-groups per treatment: LH subgroup: used to provide fertility and longevity data; RNA-1 sub-group: used to provide gene expression data following 10% adult mortality; RNA-2 sub-group used to provide gene expression data following 60% adult mortality; contingency sub-group: used to replace females that died during initial set-up and RNA females accidently lost during handling. Red horizontal arrows represent the relative longevities of each sub-group before the last individual died/was sampled. Final sample sizes were: L: LH (N=56), RNA-1 (N=54), RNA-2 (N=120); M: LH (N=45), RNA-1 (N=50), RNA-2 (N=92); H: LH (N=56), RNA-1 (N=54), RNA-2 (N=120). See Methods for full details.

To do this, we reared *D. melanogaster* larvae on three treatment diets: 20% (L), 100% (M), and 120% (H) standard Sugar Yeast Agar (SYA), and then reared the emerging adults on a 110% SYA diet. In these diets, the amount of yeast varied (20 g (L), 100 g (M), 110 g (adult) or 120 g (H) Brewer’s yeast) but other components remained the same (50 g sucrose, 15 g agar, 30 mL Nipagin, 3 mL Propionic acid per litre)^47^. We selected these three treatment diets because 20% SYA is an above-starvation but stressful diet (hence a low-quality diet), 100% SYA is the standard diet on which the *D. melanogaster* source populations were maintained (hence a medium-quality diet), and 120% SYA provides elevated yeast without overfeeding (hence a high-quality diet)^33,47,48^. We selected 110% SYA as the common garden adult diet to ensure that adults in all three treatments experienced a good-quality diet and a change to a diet different from the one experienced during larval development^45^. All flies were kept in the following conditions: 25°C, 50% humidity, 12:12 hour light:dark cycle. A pilot test showed that larvae reared on the L diet took 144 hours longer to develop into adults than did those reared on either the M or H diets (data not shown). Therefore, for the main experiment, we initiated cultures on the 20% SYA diet 144 hours before those on the 100% and 120% SYA diets, to ensure adults in all three treatments would eclose at approximately the same time^45^.

We generated larvae for the experiment from eggs collected on yeasted red grape agar oviposition plates (275 ml H20, 12.5g agar, 250 ml red grape juice, 10.5 ml of 10% w/v Nipagin solution) from four populations of wild-type Dahomey *D. melanogaster* that had been maintained on 100% SYA for at least 20 years and were derived from individuals first collected in the 1970s in Dahomey (Benin). We exposed oviposition plates to the populations for two hours to collect eggs laid within a narrow time window and thus minimise differences in development timings between individuals. After 24 hours we transferred 1^st^ instar larvae into 24 (L treatment) or 25 (M and H treatments) glass vials (vial dimensions: 75 × 25 mm, each containing 7 ml SYA; 100 larvae per vial; 2,400 larvae in total in L treatment, 2,500 larvae in total in each of the M and L treatments, hence 7,400 larvae across all treatments). We also reared a second cohort of 2,000 larvae (in 20 110% SYA vials) to produce fresh (2-3 day old) males (henceforth referred to as ‘mating males’) to provide matings with a subset of 1-2 day old females (henceforth referred to as ‘focal females’) generated by the three treatments. After this date, every eight days, we cultured enough fresh mating males to provide matings with all the surviving focal females.

We allowed larvae in the treatment vials to develop and recorded development duration as time to pupation and time to adult eclosion. We also counted numbers of larvae pupating and adults (males and females) eclosing in each vial and recorded development success as pupation success (proportion of larvae becoming pupae) and eclosion success (proportion of larvae becoming adults). (One vial from the H treatment was damaged during counting and was therefore excluded from the experiment.) To obtain focal females for the experiment, we first collected virgin adult females (i.e., females collected within four hours of eclosion, before they had reached sexual maturity, using ice anaesthesia) from the treatment vials every four hours for two days and then placed them into 110% SYA vials in groups of ten (approximately 300 virgin females in 30 vials from each treatment). At the same time, we collected approximately 900 mating males from the second cohort of flies and placed them into 90 110% SYA vials in groups of ten. To mate the focal females, we added 10 mating males into each vial of 10 focal females and allowed them to mate for 24 hours, before discarding the mating males. We then randomly selected 264 focal females from each treatment (792 focal females in total) and placed each singly into an individual vial of 110% SYA. This day was designated day 1 of the experiment. For the next seven days, we transferred each focal female to a fresh vial of 110% SYA every 24 hours. After this point, we transferred each surviving focal female to a fresh vial every 48 hours until all females were dead. All focal females were given additional opportunities to mate by, every eight days (henceforth, an ‘eight-day cycle’), placing each into a fresh vial containing a male for 48 hours. The mating males were discarded after each focal female was transferred to a fresh vial.

Within each treatment, females were each assigned a randomly generated number to allocate them to one of four sub-groups: ‘life-history’ (n = 56); ‘RNA-1’ (n = 54); ‘RNA-2’ (n = 120); and ‘contingency’ (n = 34) (Figure 1). Across all treatments, we followed 168 (3 treatments × 56 females) individual life-history females. To allow blinded observations, female treatment identity was anonymised by using a randomly generated identifier (between 1-168).

From day 1 to day 3, some focal females died in each of the three sub-groups across all treatments (L: n = 22; M: n = 85; H: n = 26). We replaced these dead females with contingency females. As the deaths stabilised from day 4, for all later analyses we used only life-history data collected from this day onwards^49^. We continued to use the contingency females to replace females in the RNA-1 and RNA-2 sub-groups that were accidentally lost or injured over the rest of the experiment. Following these procedures, final sample sizes for M females (starting on day 4) were as follows: life-history (n = 45), RNA-1 (n = 50), and RNA-2 (n = 92). The sample sizes for each sub-group in the L and H treatments on day 4 remained as stated above.

### Aim 1: Effect of larval diet treatments on fecundity-longevity relationships

To address the first aim, we used the 157 (56 L + 45 M + 56 H) life-history females to determine the effects of treatment on longevity and fertility from day 4 (Figure 1). Twice each day for the duration of the experiment, between 09:30-11:00 and 17:00-19:00, we recorded whether females remaining in the experiment were alive or dead. Once a female was found dead, we removed her and, as an index of body size, measured her thorax size (distance (mm) from the anterior margin of the thorax to the posterior tip of the scutellum, as laterally viewed, with the dorsal side upwards) using a Zeiss Discovery v12 Stereo microscope (Zeiss, Oberkochen, Germany) with Axiovision software (Zeiss). To measure fertility of females, we counted the eggs left in the previous vial each time we transferred females to fresh vials. We then allowed the eggs to develop into adult offspring over the next 11 days. In this way, for each female, we obtained egg counts and adult offspring counts every 24 hours over the first eight-day cycle (i.e., the eight days following the first mating), and every 48 hours over subsequent eight-day cycles. We froze the vials at −4°C and counted all adult offspring following the death of the last focal female (88 days post-eclosion).

### Aim 2: Effect of larval diet treatments on gene expression in adult females

To address the second aim, we used the RNA-1 and RNA-2 females to produce samples for gene expression profiling by mRNA-seq (Figure 1). For both sub-groups within each treatment, we recorded mortality in the same way as for the life-history females. Within each treatment, we collected surviving RNA-1 females after 10% of the total RNA-1 females had died (5/54 mortalities in the L and H treatments; 5/50 mortalities in the M treatment). We collected surviving RNA-2 females after 60% of the total RNA-2 females had died (72/120 mortalities in the L and H treatments; 55/92 mortalities in the M treatment). These time-points (10% and 60%) were selected to represent low and high mortality thresholds, respectively. We chose time-points based on relative (rather than absolute) time to account for differences in ageing between different treatments and to facilitate comparisons with other studies^39,43^. To sample females at each time-point, we waited until the requisite number of deaths had been reached, and then anaesthetised the remaining females within the sub-group (within the treatment) on ice at 10:30 the following morning. We then froze all of the anaesthetised females in liquid Nitrogen and stored them at −80°C. We continued this procedure until every female had died or been sampled. Overall, we collected and stored the following numbers of females from each treatment **L:** RNA-1: N = 49 (out of 54), RNA-2: N = 48 (out of 120); **M:** RNA-1: 45 (out of 50), RNA-2: 37 (out of 92); **H:** RNA-1: N = 49 (out of 54), RNA-2: N = 48 (out of 120).

### Dissections and RNA extraction for RNA-sequencing

Analyses of the life history females showed that M and H treatment females had dissimilar fecundity-longevity relationships (see Results). We therefore sequenced the RNA-1 and RNA-2 mRNA samples from these two treatments alone (and not from the L treatment). We dissected females on a glass Petri dish over dry ice. For each female, we collected the head, fat body, and ovaries, and we then pooled each tissue across females into samples of 11-12 individuals each, with 3-4 such samples being created per sub-group within each treatment. The fat body samples were collected from each isolated abdomen by first removing the ovaries (which were sampled separately) and gut (which was discarded), and then pooling the remains of the abdomen. We disrupted the tissue samples with a micropestle and mixed in an appropriate volume of Tri-reagent (100 μl per 10 mg of tissue; Sigma-Aldrich, Gillingham, Dorset, UK). We then extracted RNA using the Direct-zol™ RNA extraction kit (Zymo Research, Irvine, CA, USA) according to the manufacturer’s protocol, and carried out an additional Dnase treatment using the Turbo™ DNA-free kit (Thermo Fisher Scientific, Loughborough, UK) again according to the manufacturer’s protocol. Following these procedures, we produced the following samples: M RNA samples (three samples of 12 heads, three samples of 12 fat bodies, three samples of 12 ovaries from each of the RNA-1 and RNA-2 sub-groups, generating 3 samples of 12 pooled individuals each × 3 tissues × 2 subgroups = 18 RNA biological replicates in total); and H RNA samples (four samples of 11-12 heads, four samples of 12 ovaries, and four samples of 12 abdomens from each of the RNA-1 and RNA-2 sub-groups, generating 4 samples of 11-12 pooled individuals each × 3 tissues × 2 sub-groups = 24 RNA biological replicates in total). Of the collected samples, to equalise sample sizes, we sent three RNA samples of each treatment, tissue, and sub-group to Edinburgh Genomics (Edinburgh, UK) for Illumina 100 base pair, paired-end sequencing on a NovaSeq6000 sequencer (18 biological replicates/treatment × 2 treatments = 36 RNA samples in total).

## Statistical analysis

All statistical analyses were conducted with the R (version 4.0.3)^50^ statistical programming platform in RStudio, using the ‘lme4 (v1.1-28)’^51^, ‘glmmTMB (v1.1.3)’^52^, and ‘survival (v3.4-0)’^53^ packages. To test whether models of count data were overdispersed we used the ‘dispersiontest’ function in the ‘AER (v1.2-9)’^54^ package. We determined the significance of the fixed effects using a likelihood ratio test.

### Effect of larval diet treatments on fecundity-longevity relationships

#### Development duration, development success, and adult female thorax size of larvae reared on L, M, and H larval diets

We first investigated whether treatment affected development duration, development success, and adult female thorax size, as further described in the supplementary information.

#### Longevity of females reared on L, M, and H larval diets

We investigated the effect of larval diet on longevity using Cox Proportional Hazards regression analyses. Longevity for each female was calculated as the number of days between the date the female eclosed and the date of the female’s death. We fitted Cox models using the ‘coxph’ function from the ‘survival’ package in R. For each analysis, the longevity data satisfied the coxph function’s assumptions of proportional hazards (the relationship between the scaled Schoenfeld residuals and time was zero) and there was a linear relationship between the log hazard and its covariates. We first modelled longevity across the whole experiment and treated all females that that escaped or were accidently injured during transfers as censors (5/157 life-history females across all three treatments). Differences in survival appeared to converge after day 55 (Figure 2), prompting us to conduct additional analyses with the data split into two groups, i.e., females that died before day 55 (‘early-death females’) or after day 55 (‘late-death females’). In each case, there was no effect on the significance of treatment as a predictor, so only the model outputs from across the whole experiment are shown in the results. The initial maximal model included the main effect (treatment), an additional fixed effect (adult female thorax size), and an interaction term (treatment:adult female thorax size). Since the model with the lowest AICc value did not include adult female thorax size or the interaction term, these factors were dropped from the final model.

**Figure 2.**
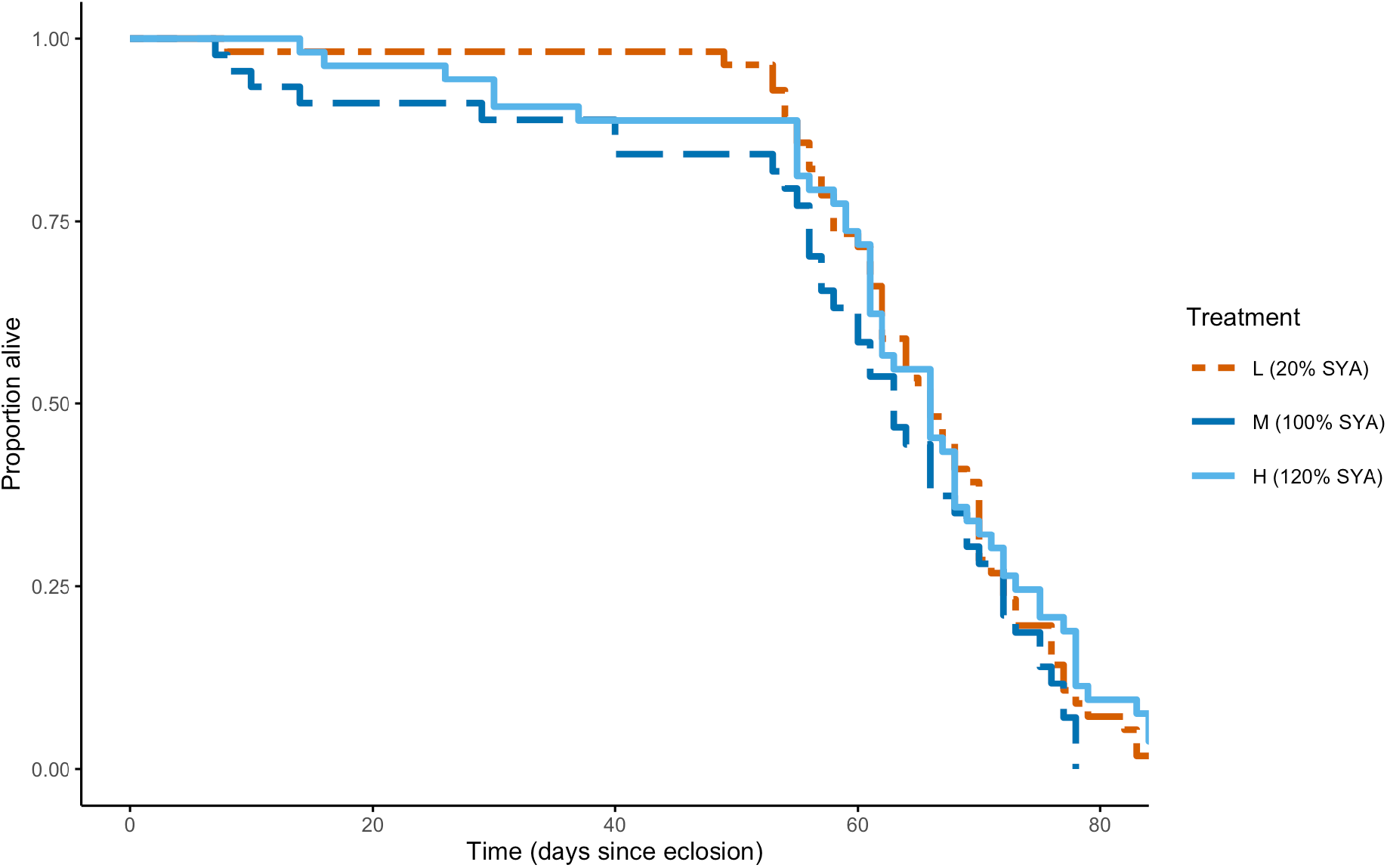
Proportion alive against time (days since eclosion) for adult female *Drosophila melanogaster* reared on L (20% SYA, orange dashed line, n = 56), M (100% SYA, dark blue dashed line, n = 45), H (120% SYA, light blue solid line, n = 56) treatment diets as larvae, and on a 110% SYA diet as adults. There was no significant effect of treatment on proportion alive over time, and hence no effect on female longevity.

#### Fertility of females reared on L, M, and H larval diets

To assess the effect of treatment on fertility we analysed four time-dependent, i.e repeated, measures of fertility for each life-history female: 1) early-life egg production: the number of eggs produced every 24 hours for the first eight-day cycle for each female; 2) early-life offspring production: the number of adult offspring (i.e., eggs that reached adulthood) produced every 24 hours for the first eight-day cycle for each female; 3) whole-life egg production: the number of eggs produced during each eight-day cycle following mating across the whole life of each female (hence, within each eight-day cycle, summed across all eight days that eggs were counted for the first eight-day cycle, and across all four days that eggs were counted for each subsequent eight-day cycle); 4) whole-life offspring production: the number of adult offspring produced during each eight-day cycle following mating across the whole life of each female (within each eight-day cycle, adult offspring counts summed as for whole-life egg production). We tested for an effect of treatment and age on each of the four measures using generalised linear mixed effects models (glmm) to account for repeated measures on the same females. In each case the data were found to be significantly overdispersed and significantly zero-inflated. To account for this, we fitted zero-inflated glmms with negative binomial error structure using the ‘glmmTMB’ function from the glmmTMB R package^52^. Due to the non-linear effect of age on egg and offspring production we fitted each model with age as a quadratic term. For early-life egg production and early-life offspring production models, treatment, age (in days), and their interaction terms were fitted as fixed effects, and a unique identifier for each female was fitted as a random effect. For whole-life egg production and whole-life offspring production, treatment, age (in eight-day cycles), treatment - age interaction terms and adult female thorax size were fitted as fixed effects, and a unique identifier for each female was fitted as a random effect. We then compared each final model with its simplified model equivalents, and the results using the model with the lowest AICc value (which retained treatment as a fixed factor) are reported below. Following this, the models with the lowest AICc values did not include adult female thorax size, so this factor was dropped from all final models. The model rankings (from lowest to highest AICc value) are reported in Table S1.

In addition to the four time-dependent measures of fertility, we also tested for an effect of treatment on fertility using two additional lifetime measures for each female: 1) total eggs produced: the total number of eggs produced (summed across all eight-day cycles); 2) total offspring produced: the total number of adult offspring produced (summed across all eight-day cycles). We used glms with negative binomial error structure to test for an effect of treatment on each of these measures. We then determined the significance of the predictor effect by comparing the glm with an intercept-only glm using a likelihood ratio test.

We also assessed the effect of treatment on egg viability for each female (the total number of adults developing from eggs as a proportion of the total number of eggs that were produced during each eight-day cycle). We analysed these proportion data using a glmm (with binomial error distribution) using the ‘glmer’ function in the ‘lme4’ package. Egg viability was the proportional response variable. Treatment, age (in eight-day cycles), and their interaction terms were fitted as fixed effects, and a unique identifier for each female was fitted as a random effect. We then used model simplification as described above. The model rankings (from lowest to highest AICc value) are reported in Table S1.

#### Fecundity-longevity relationships of females reared on L, M, and H larval diets

To test the hypothesis that the directionality of the fecundity-longevity relationship in adult females becomes less negative when females are reared as larvae on higher quality diets, we used an ANCOVA test to determine the combined effects of diet and longevity on mean egg and mean offspring production for each female (respectively, the mean number of eggs or adult offspring produced every day over each female’s lifetime). Mean egg and offspring production were calculated for each female by dividing total eggs produced (or total offspring produced) by longevity. For the L treatment, a test using Cook’s distance showed that one female was an outlier (ID: life-history-47), so we also assessed the impact on the relationship when life-history-47 was removed. This was found to have no effect on the overall result, so the results with life-history-47 included are reported. Adult female thorax size was included as an additional fixed effect. The model with the lowest AICc value did not include adult female thorax size so this factor was then dropped from the final model. Following this analysis, we hypothesised that the observed fecundity-longevity relationships in each treatment were partly driven by early-death females (see Results); to test this, we repeated the analysis omitting all early-death females (including life-history-47).

We used these ANCOVA analyses to determine which of the three treatments would show the strongest contrast in fecundity-longevity relationships, and thus which treatments to sequence for mRNA-seq analysis. Following analysis, we determined that the M and H treatments showed the greatest contrast in the directionality of the fecundity-longevity relationship (see Results).

### Bioinformatic analysis

#### Quality assessment of mRNA-seq reads

We used FastQC v0.11.9^55^ to examine base quality and potential adapter contamination in each sample, and combined the results for each sample into a report for each tissue (head, fat body, and ovaries) using the MultiQC v1.9 Python library^56^ with Python v3.7^57^ (Supplementary files S1-3). We then aligned reads against the *D. melanogaster* genome (Drosophila_melanogaster.BDGP6.22.dna.toplevel.fa) using HISAT2 v2.1.0^58^ and recorded mapping statistics (Table S2). We used the HISAT2 alignment files to assess gene body coverage and junction saturation using the rSeQC v3.0.1 Python library^59^ with Python v3.7^57^. We determined that each mRNA-seq library passed quality assessment and therefore retained all libraries in the analysis.

### Effect of larval diet treatments on gene expression in adult females

#### Age-related gene expression

We pseudoaligned reads to the *D. melanogaster* transcriptome (Drosophila_melanogaster.BDGP6.22.cdna.all.fa) with Kallisto v0.46.1^60^ and we used the tximport package v1.16.1 (Soneson et al. 2016) in R (v4.0.1)^50^ to estimate transcript counts for each gene. We used these estimated counts for differential expression analysis in R (v4.0.1)^50^ with the DESeq2 package v1.28.1^61^ using an FDR adjusted *p*-value threshold of 0.05 and the model ~ condition where condition was a categorical factor denoting the combined treatment and time point of a sample. We produced boxplots of the normalised count data and principal component analysis from DESeq2 for each tissue to check normalisation and library clustering, respectively (Figures S1 – S3). We generated four lists of differentially expressed genes (DEGs) for each tissue: genes more highly expressed in RNA-2 than RNA-1 (up-regulated genes) and genes more highly expressed in RNA-1 than RNA-2 (down-regulated genes) for both M and H treatments (Table S3 - S5)).

We performed Fisher’s Exact Tests to detect significant overlaps between each pair of lists from M and H in each tissue (head, fat body, and ovaries) and each direction of differential expression with respect to age (up- and down-regulated with age), resulting in 6 comparisons in total. For this, we used custom R (v4.0.1) scripts^50^ and we used Bonferroni correction to account for multiple testing.

#### Gene Ontology

We performed GO enrichment analysis in R (v4.0.1)^50^ via the clusterProfiler package (v3.16.1)^62^ using biological processes GO annotations from the org.Dm.eg.db package (v3.11.4)^63^. We used the over-representation test^64^ to identify GO terms that were significantly overrepresented (*p* < 0.05 after adjustment for multiple testing with *Benjamini-Hochberg*) in a set of DEGs against a background consisting of all genes that were expressed in the relevant tissue. We reduced redundancy in the resulting enriched GO terms using the GoSemSim package (v2.14.2)^65^.

#### Comparisons with age- and/or ageing-related genes from other studies

To determine whether a change in the directionality of the fecundity-longevity relationship altered the pattern of gene-expression for age-related genes previously isolated in *D. melanogaster*, we identified two *D. melanogaster* mRNA-seq studies^66,67^ in which tissues (fat bodies^66^; brains in^67^) and relative mortality at sampling were comparable to those in the current study. For these studies^66,67^, we downloaded the total gene list and the list of genes that were differentially expressed (as defined by each study) between the appropriate time points. For Pacifico et al.^67^, we used genes differentially expressed between 5 day old and 30 day old adults, as the percentages of the cohort surviving were broadly similar (5 days: ~30% mortality, 30 days: ~55% mortality) to those in the current study. We also compared the DEGs from the current study to the *D. melanogaster* genes in the GenAge database^68^, which contains experimentally validated genes associated with ageing. To determine whether components of the IIS-TOR network were differentially expressed with age depending on the directionality of the fecundity-longevity relationship in *D. melanogaster*, we identified the relevant *D. melanogaster* genes associated with the TI-J-LiFe network^32^.

In these analyses, we used Fisher’s Exact Tests to detect significant overlaps between lists of DEGs from M and H and the other lists of insect genes. We compared the DEGs from Pacifico et al.^67^ and Chen et al.^66^ to the DEGs from M and H in head and fat body (respectively) in the current study, comparing gene lists that were differentially expressed in same manner with respect to time (total of 8 comparisons). We compared the GenAge gene list to the combined up- and down-regulated genes within each tissue and treatment in the current study, as we had no expectation as to whether the GenAge genes would be up- or down-regulated with age (total of 6 comparisons). We modified the approach taken by Korb et al.^32^ to compare the DEGs from the current study to the lists of *D. melanogaster* genes associated with the TI-J-LiFe network. We combined up- and down-regulated DEGs within each tissue and treatment in the current study and ranked them by log fold change in expression with time. We then extracted the 50 genes with the most positive log fold change and the 50 genes with the most negative log fold change within each tissue/treatment (hereafter, top ± 50 genes). Likewise, we extracted the top ± 100 genes, top ± 200 genes, top ± 300 genes, and top ± 500 genes. We then compared the TI-J-LiFe network gene list with the top ± genes lists for each tissue/treatment combination (total of 36 comparisons). We performed Fisher’s Exact Tests using R (v4.0.1)^50^ and accounted for multiple testing using Bonferroni correction.

## Results

### Effect of larval diet treatments on fecundity-longevity relationships

#### Development duration, development success, and adult female thorax size of larvae reared on L, M, and H larval diets

Larvae reared on the L treatment had significantly increased development durations and significantly decreased adult female thorax sizes compared to the other two treatments but there was no effect of treatment on development success (Figures S5 – S9; see Supplementary Results for details). Larvae reared on the M treatment also had significantly increased development durations compared to larvae reared on the H treatment (Figures S5 – S6; see Supplementary Results for details). Larvae reared on the M and H treatments had significantly increased adult female thorax sizes compared to larvae reared on the L treatment, but were not significantly different from each other (Figures S5 – S6; see Supplementary Results for details).

#### Longevity of females reared on L, M, and H larval diets

The longevities of L females (mean longevity (SD) = 65.2 (16.3) days), M females (mean longevity (SD) = 58.7 (19.1) days) and H females (mean longevity (SD) = 63.3 (11.7) days) did not differ significantly (coxph: M > L: z = −1.234, p = 0.217; M > H: z = −1.799, p = 0.072; Figure 2). Therefore, there was no effect of the treatments on adult female longevity.

#### Fertility of females reared on L, M, and H larval diets

H females had significantly higher early-life egg production than M and L females (mean eggs produced per day (SD): H = 12 (7.9), M = 10.9 (8.5), L = 9.6 (7.3); glmm: z = 1.999, p = 0.046; Figure 3a, Tables S1, S6). However, there was no effect of treatment on whole-life egg production (glmm: H > M: z = 1.300, p = 0.194; M > L: z = 0.083, p = 0.934; Figure 3a, Tables S1, S6) or total eggs produced (LRT: df = 2, χ^2^ = 0.221, p = 0.895; Figure 3b). Despite the early-life differences in egg production, there were no significant differences between treatments in early-life offspring production (glmm: H > M: z = 1.227, p = 0.220; L > M: z = 0.206, p = 0.837), whole-life offspring production (glmm: H > M: z = 0.162, p = 0.871; L > M: z = 0.745, p = 0.474; Figure 3c, Tables S1, S6), or total offspring produced (LRT: df = 2, χ^2^ = 0.008, p = 0.996, Figure 3d). There were also no significant differences between treatments in egg viability (glmm: H > M: z = −0.578, p = 0.563; L > M: z = −0.030, p = 0.976; Figure S4, Tables S1, S6). In all three treatments, whole-life egg production, whole-life offspring production, and egg viability significantly decreased over time (glmm: whole-life egg production: z = −35.994, p <0.001; whole-life offspring production: z = −21.419 p <0.001; egg viability: z = −14.067, p < 0.001; Figures 3a, 3c, S4, Tables S1, S6) and there were no interactions between treatment and time for any of these variables (Tables S1, S6).

**Figure 3.**
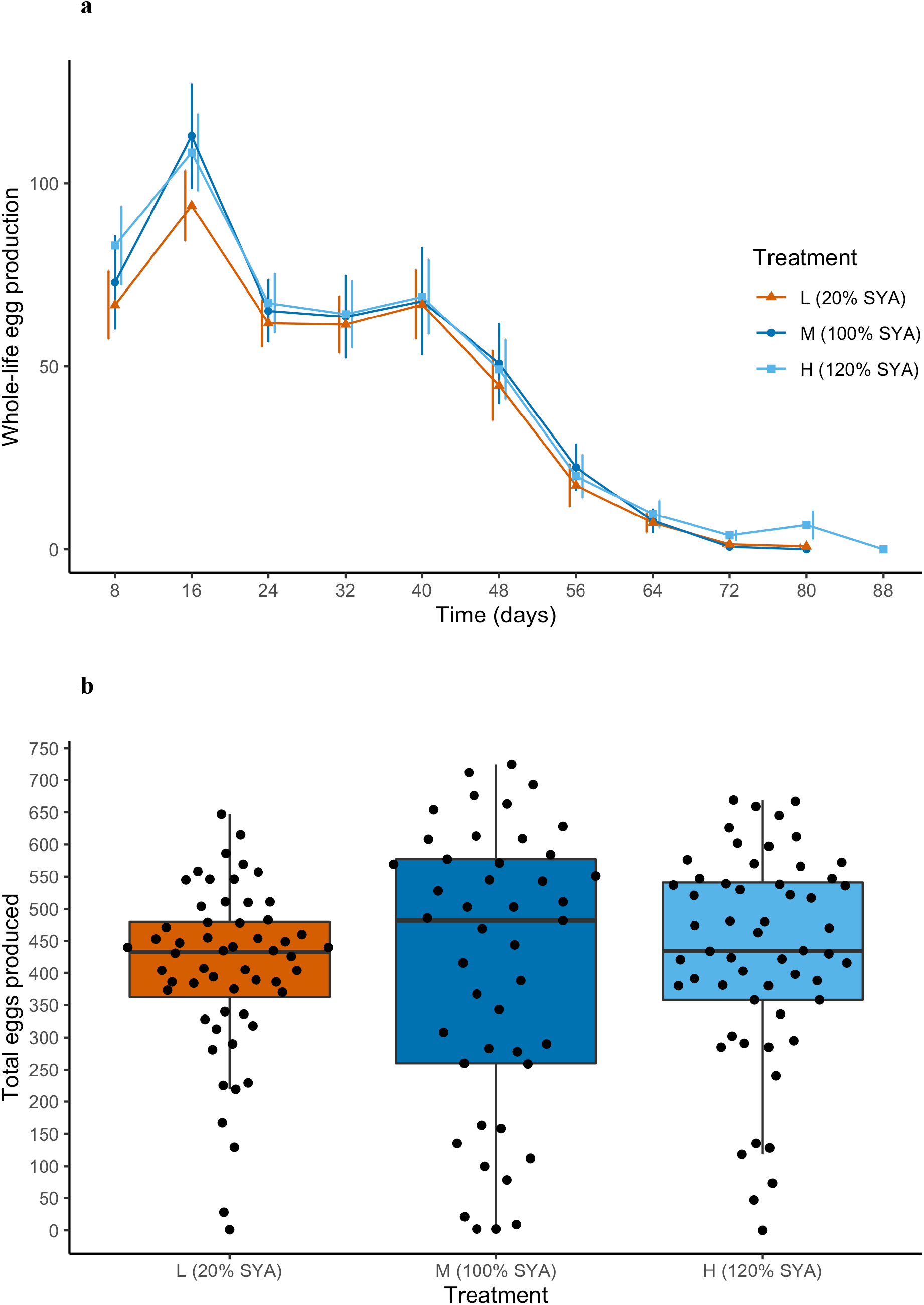
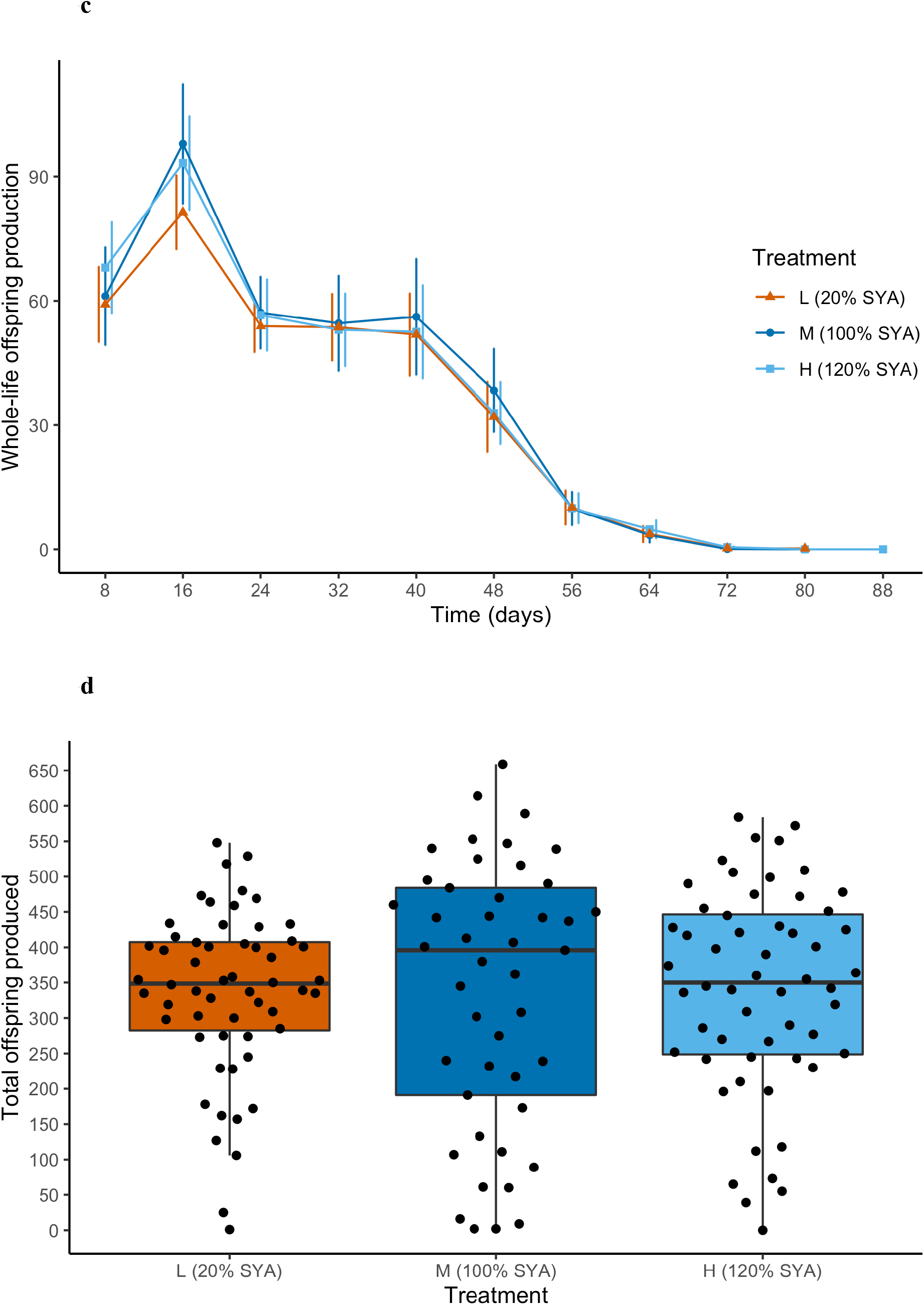
Fertility (egg and adult offspring production) in adult female *Drosophila melanogaster* reared on L (20% SYA, orange, n = 56), M (100% SYA, dark blue, n = 45), H (120% SYA,light blue, n = 56) treatment diets as larvae, and on 110% SYA diet as adults. A) Whole-life egg production (number of eggs produced per eight-day cycle) as a function of time (represented by the last day of each successive eight-day cycle) for L (triangles), M (circles), H (squares) females. Error bars: 1 S.D. Although whole-life egg production did not differ between treatments, early-life egg production (daily egg production during the first eight-day cycle) was significantly higher for H females compared to M and L females. For all treatments, whole-life egg production declined significantly with time. B) Total eggs produced (sum total number of eggs produced per female, black circles). Horizontal bars: median (across all females within each treatment); boxes: interquartile ranges; whiskers: ranges up to 1.5 × the interquartile range. There was no significant difference in total eggs produced per female between treatments. C) Whole-life offspring production (number of adult offspring produced per eight-day cycle) as a function of time (represented by the last day of each successive eight-day cycle) for L (triangles), M (circles), H (squares) females. Error bars: 1 S.D. Neither early-life offspring production (daily offspring production during the first eight-day cycle) or whole-life offspring production were significantly affected by treatment. For all treatments, whole-life offspring production decreased significantly with time. D) Total offspring produced (sum total number of eggs produced per female, black circles). Horizontal bars: median (across all females within each treatment); boxes: interquartile ranges; whiskers: ranges up to 1.5 × the interquartile range. There was no significant difference in the total offspring produced per female between treatments.

#### Fecundity-longevity relationships of females reared on L, M, and H larval diets

The relationship between longevity and mean fertility (measured as either mean daily egg production or mean daily offspring production) showed a significant interaction with treatment (ANCOVA, Eggs: F = 11.991, df = 2, 146, p < 0.001; Offspring: F = 9.427, df = 2, 146, p < 0.001; Figure 4). L females showed no significant relationship between longevity and mean fertility (linear regression, eggs: F = 1.619, df = 1, 54, p = 0.209, R^2^ = 0.011; offspring: F = 1.096, df = 1, 54, p = 0.299, R^2^ = 0.002), M females a significant positive relationship (eggs: linear regression, F = 14.63, df = 1, 41, p < 0.001, R^2^ = 0.245; offspring: F = 12, df = 1, 41, p = 0.001, R^2^ = 0.208) and H females a significant negative relationship (eggs: F = 7.664, df = 1, 51, p = 0.008, R^2^ = 0.114; offspring: F = 5.529, df = 1, 51, p = 0.023, R^2^ = 0.08) (Figure 4). When early-death females (females dying before day 55) were omitted from the analysis, there was still a significant interaction between longevity, fertility, and treatment (ANCOVA, Eggs: F = 4.409, df = 2, 125, p = 0.014; Offspring: F = 3.463, df = 2, 125, p = 0.034). L females still showed no significant relationship between longevity and mean fertility (eggs: F = 1.255, df = 1, 48, p = 0.268, R^2^ = 0.005; offspring: F = 1.516, df = 1, 48, p = 0.224, R^2^ = 0.01) and H females still showed a significantly negative relationship (eggs: F = 12.03, df = 1, 45, p = 0.001, R^2^ = 0.193; offspring: F = 5.057, df = 1, 45, p = 0.029, R^2^ = 0.081). However, M females showed no significant relationship between longevity and mean fertility (eggs: F = 1.132, df = 1, 32, p = 0.261, R^2^ = 0.009; offspring: F = 2.032, df = 1, 32, p = 0.164, R^2^ = 0.03). Nonetheless, M and H females therefore still differed in the directionality of their fecundity-longevity relationships.

**Figure 4.**
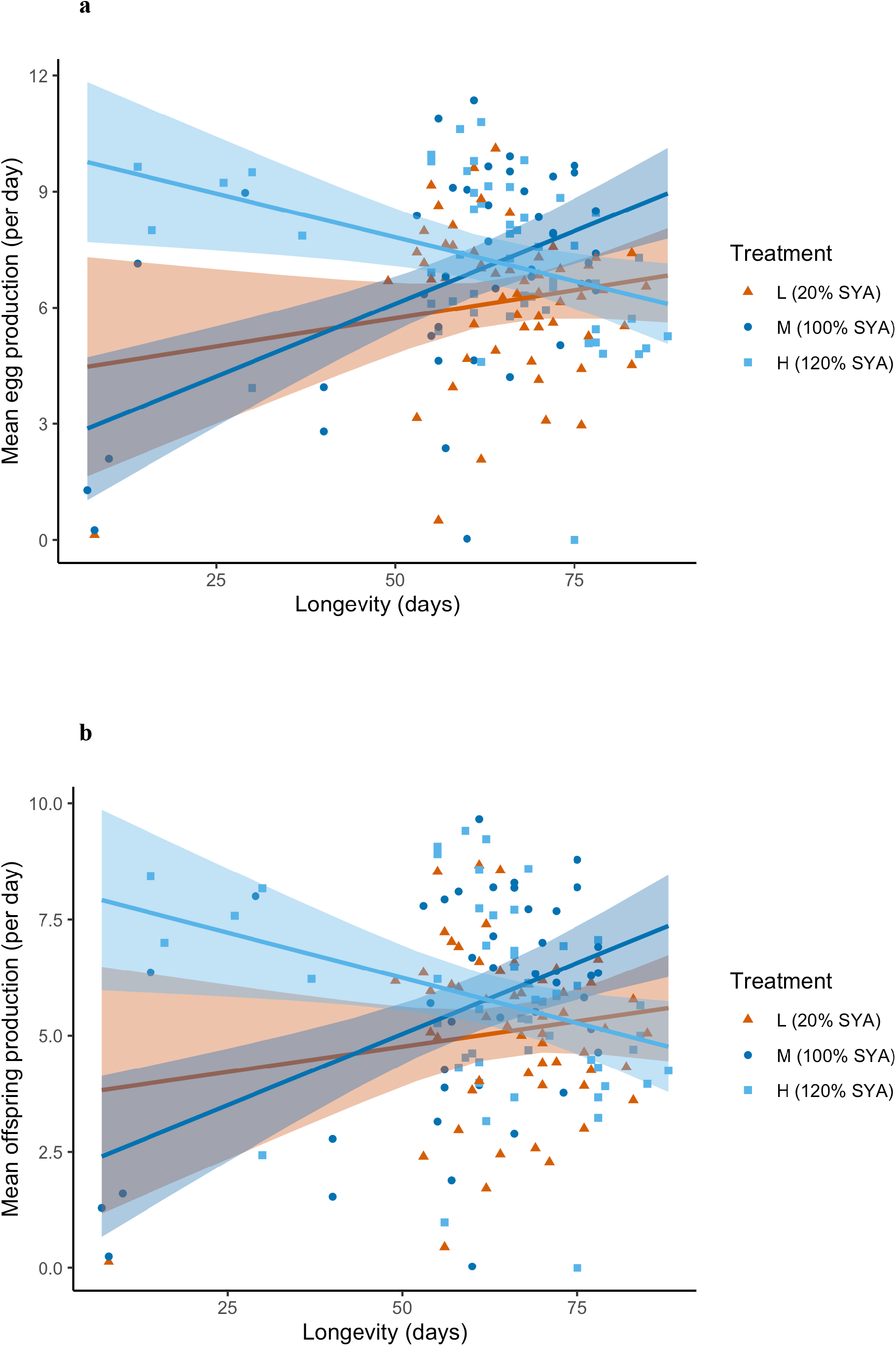
Fecundity-longevity relationships in adult female *Drosophila melanogaster* reared on L (20% SYA diet, orange, n = 56), M (100% SYA diet, dark blue, n = 45), H (120% SYA diet, light blue, n = 56) treatment diets as larvae, and on 110% SYA diet as adults. A) Longevity versus mean egg production (mean number of eggs produced every day over each female’s lifetime). B) Longevity versus mean offspring production (mean number of adult offspring produced every day over each female’s lifetime). The solid line and shading represent each linear relationship and set of confidence intervals for each treatment, respectively. For both mean egg production and mean offspring production, females on the 20% SYA diet showed no significant fecundity-longevity relationship, on the 100% SYA diet showed a significant positive relationship, and on the 120% diet showed a significant negative relationship.

These results support the hypothesis that the directionality of the fecundity-longevity in adult female *D. melanogaster* can be changed by larval diet. However, the signs of these relationships were not the same as those we predicted for their respective diets (being of opposite signs in the analysis which included early-death females), and therefore the contrast in directionality was also not the same as the one hypothesised.

### Effect of larval diet treatments on gene expression in adult females

#### Age-related gene expression

Sequencing (by mRNA-seq) of the 36 libraries resulted in a mean of 71,545,047 read pairs per library for head, 72,699,206 read pairs per library for fat body and 85,320,432 reads pairs per library for ovaries (Table S7). The libraries pseudoaligned to the *D. melanogaster* transcriptome with a mean 89.9% match (range 87.4 – 92.1%) for head, mean 92.0% match (91.3 – 92.9%) for fat body, and mean 92.3% match (89.7 – 94.4%) for ovaries (Table S8).

In total, across both treatments from which adult females were sequenced (M and H larval diets) and both time-points (RNA-1 and RNA-2), there were 6,797 differentially expressed genes (DEGs) in head, 5,702 DEGs in fat body and 7,320 DEGs in ovaries. Within tissues and treatments, similar numbers of genes were up- and down-regulated in comparisons across the time points. In head, in M females, 2,006 genes were more expressed in RNA-2 than RNA-1 (up-regulated genes) and 2,125 genes were more expressed in RNA-1 than RNA-2 (down-regulated genes); and in H females, 1,425 genes were up-regulated and 1,241 genes were down-regulated (Figure S10, Table S3). In fat body, in M females, 1,555 genes were up-regulated and 1,275 genes were down-regulated; and in H females, 1,395 genes were up-regulated and 1,477 genes were down-regulated (Figure S11, Table S4). In ovaries, in M females, 1,247 genes were up-regulated and 1,329 genes were down-regulated; and in H females, 2,478 genes were up-regulated and 2,266 genes were down-regulated (Figure S12, Table S5).

We compared lists of M and H treatment DEGs within each tissue to reveal whether the gene expression profiles were similar. In head, 41.4% of up-regulated M DEGs were shared with up-regulated H DEGs, and 37.4% of down-regulated M DEGs were shared with down-regulated H DEGs. In fat bodies, 35.0% of up-regulated M DEGs were shared with up-regulated H DEGs, and 35.7% of down-regulated M DEGs were shared with down-regulated H DEGs. In ovaries, 80.4% of up-regulated M DEGs were shared with up-regulated H DEGs, and 77.5% of down-regulated M DEGs were shared with down-regulated H DEGs. Overall, all comparisons (6/6) of M and H DEGs showed significant overlaps (mean percentage overlap relative to the M treatment = 51.23%; Fisher’s Exact Test,*p* > 1 x 10^-124^ in all cases) (Figure 5, Tables S9 and S10).

**Figure 5.**
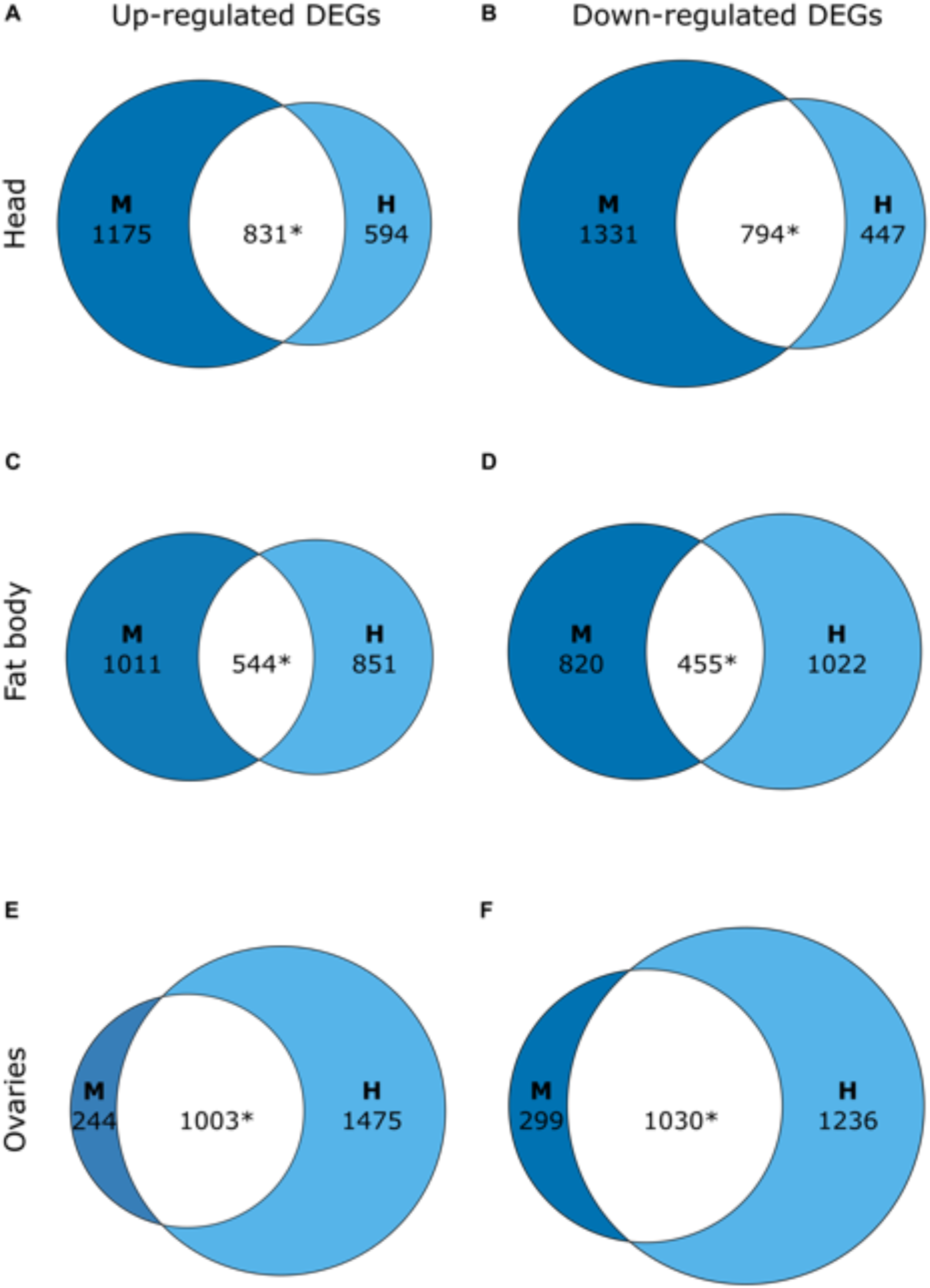
Comparison of changes in gene expression profiles with time in three tissues between M (100% SYA) and H (120% SYA) adult female *Drosophila melanogaster* as determined by mRNA-seq. Euler diagrams of overlaps between differentially expressed genes (DEGs), i.e. genes differentially expressed between RNA-1 females and RNA-2 females and shared between M females (dark blue circles) and H females (light blue circles) for: A), B), head; C), D), fat body; and E), F), ovaries. Asterisks (*) denote significant overlap in DEGs (Fisher’s exact test, *p* < 0.05 after Bonferroni correction). Up-regulated DEGs: DEGs significantly more expressed in RNA-2 females than RNA-1 females, i.e., that increase expression with age; down-regulated DEGs: DEGs significantly more expressed in RNA-1 females than RNA-2 females, i.e., that decrease expression with age. Results of statistical tests are in Table S9 and identities of overlapping genes are in Table S10.

Hence, there were significant similarities between treatments in age-related DEGs in all three tissues. However, head and fat body showed greater differences (less overlap) than ovaries.

#### Gene ontology

Gene ontology (GO) enrichment analysis isolated 1,323 nonredundant enriched GO terms for the DEGs across all treatments and tissues (Table S11). In head, in M and H females, up-regulated DEGs were enriched for a wide variety of processes with no obvious pattern (Table S11), and down-regulated DEGs were enriched for GO terms associated with metabolic and biosynthetic processes (29/59 and 24/63 terms, respectively) and with the eye and sensing light stimuli (8/59 and 7/63 terms, respectively) (Table S11). In head, in M females, down-regulated DEGs were also enriched for terms associated with mitochondria (7/59 terms). In H females, down-regulated DEGs were enriched for terms associated with transport (9/63) and the nervous system (5/63) (Table S11).

In fat body, in M females, up-regulated DEGs were enriched for GO terms associated with metabolic and biosynthetic processes (40/125 terms) and immune/defense response (33/125 terms), and in H females, up-regulated DEGs were enriched for GO terms associated with metabolic and biosynthetic processes (20/36) and with regulation of the cell cycle and associated nuclear and chromosomal changes (11/36 terms) (Table S11). In M females, down-regulated DEGs were enriched for a wide variety of processes with no obvious pattern (Table S11). In H females, down-regulated DEGs were enriched for GO terms associated with metabolic and biosynthetic processes (13/57 terms).

In ovaries, in M females, up-regulated DEGs were enriched for GO terms associated with a wide variety of processes with no obvious pattern, and in H females, up-regulated DEGs were enriched for GO terms associated with mitosis, meiosis and associated nuclear and chromosomal changes (22/84 terms). In M females, down-regulated DEGs were enriched for GO terms associated with metabolic and biosynthetic processes (24/44 terms) (Table S11), and in H females, down-regulated DEGs were enriched for a wide variety of processes with no obvious pattern (Table S11).

For the GO terms GO:0007568 ‘aging’ and/or GO:0010259 ‘multicellular organism aging’, we found enrichment in DEGs up-regulated in the head of M and H females, and in DEGs down-regulated in the fat body and ovaries of H females (Table S11).

Hence, GO term enrichment analysis showed that the biological functions of age-related DEGS differed between each treatment.

#### Comparisons with age- and/or ageing-related genes from other studies

We compared lists of M and H treatment DEGs to previously identified *D. melanogaster* age-related genes. For the comparison with two previous tissue-specific mRNA-seq *D. melanogaster* studies^66,67^7/8 comparisons showed significant overlap between DEGs in the current study and the previous studies (mean percentage overlap of significant comparisons = 16.14% of DEGs in the current study; Fisher’s Exact Test, *p* < 1 x 10^-6^ in each case). Only the comparison between DEGs up-regulated in H in fat body did not show significant overlap (Fisher’s Exact Test, *p* = 0.03) (Tables S12, S13). For the comparison with genes in the *D. melanogaster* GenAge database^68^, 2/6 comparisons showed significant overlap. The two significant overlaps were as follows: (a) 98/194 (50.5%) of the GenAge genes were up-regulated in head in M (Fisher’s Exact Test, *p* = 1.01 x 10^-7^); and (b) 68/194 (35.1%) of the GenAge genes were up-regulated in fat body in M (Fisher’s Exact Test, *p* = 9.46 x 10^-5^) (Tables S14, S15). Genes from the *D. melanogaster* GenAge database that were differentially expressed with age in head or fat body of the current study are shown in Figure S13.

We also compared expression of M and H DEGs to that of genes associated with the TI-J-LiFe network^32^(Figure 6). In M female heads, for 1/6 comparisons, 59/123 (48%) of the TI-J-LiFe genes significantly overlapped with M female head DEGs (Fisher’s Exact Test, *p* = 0.00003). The significant comparison occurred when all M head DEGs were included in the analysis (Table S16). However, there were no significant overlaps with TI-J-LiFe genes in any other tissue/treatment combinations (0/30 comparisons showed overlap) (Figure 5, Tables S16, S17).

**Figure 6.**
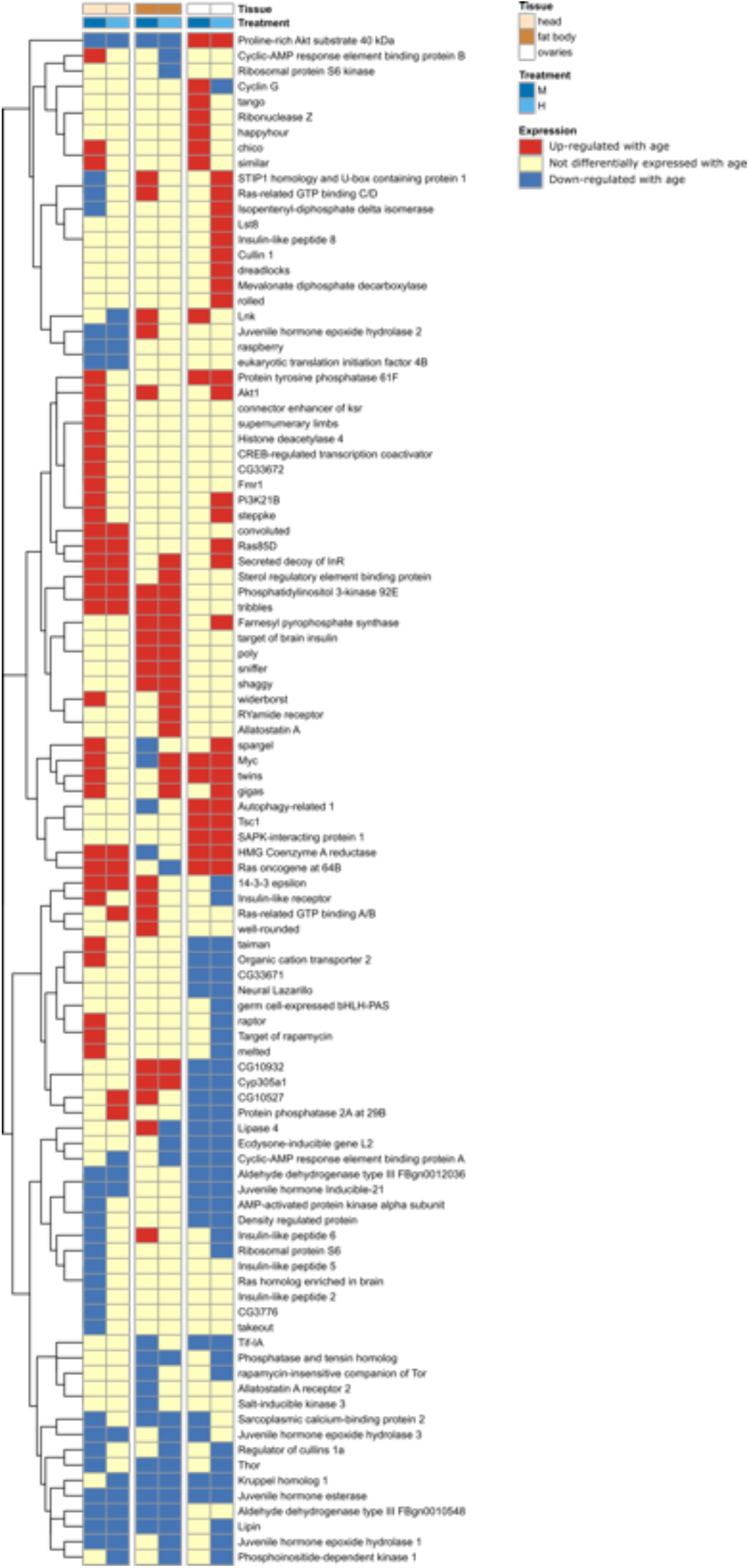
Results of comparison of age-related genes in M (100% SYA) and H (120% SYA) adult female *Drosophila melanogaster* and genes in the TI-J-LiFe network. Each row represents an individual gene from *Drosophila melanogaster* described as a component of the TI-J-LiFe network by Korb et al.^32^. Each column shows the age-related expression status of focal genes in a given treatment and tissue in *D. melanogaster* females in the current study. Vertical breaks separate the three tissues studied (head, fat body, and ovaries). The dendogram at left groups genes according to their gene expression patterns.

Overall, differences in treatments, and hence in the directionality of the fecundity-longevity relationship, were associated with both similarities and differences in age-related gene expression in adults. A significant overlap in DEGs between M and H treatments was found in all tissues examined; however, there were still many DEGs whose expression differed between treatments. For example, only the M treatment showed significant overlaps in gene expression with the GenAge and TI-J-LiFe gene lists.

## Discussion

We tested for an effect of larval diet on the directionality of the fecundity-longevity relationship and for associated age-related gene expression changes in adult *D. melanogaster* females. We found that larval diets significantly affected development duration (but not development success; Figures S2 - S4) and adult female thorax size (Figure S6), but had no effect on longevity (Figure 2) and relatively small effects on fecundity (i.e., the H diet had significantly higher early-life egg production than the other two treatments, but the effect size was modest; Figure 3a). Despite this, there was a clear effect of treatment on the directionality of the fecundity-longevity relationship (Figure 4). Specifically, L females showed no significant relationship between fertility and longevity, M females showed a significant positive relationship (or no significant relationship in the analysis omitting early-death females) and H females a significant negative relationship. Larval diet quality also had a marked effect on age-related gene expression in adult females, with M and H treatments showing significantly overlapping, but distinct, sets of age-related genes.

### Effect of larval diet treatments on fecundity-longevity relationships

Other things equal, the standard evolutionary theory of ageing predicts a negative fecundity-longevity relationship; however, there are many examples where the fecundity-longevity relationship becomes uncoupled or is positive (see Introduction). Like many other organisms, *D. melanogaster* typically exhibits a negative fecundity-longevity relationship in females (Flatt 2011), although longevity and fecundity can become uncoupled in long-lived mutants^8,9,12^ and on different quality diets^36,41^. Our results show that, in addition, the fecundity-longevity relationship can change directionality in response to changes in larval diet quality. However, this did not result in longer-lived females incurring greater costs of reproduction, as there were no differences in adult fertility or longevity between treatments. All females also showed reproductive senescence, i.e. their reproduction declined significantly with time in all treatments (Figure 3a).

These results could help explain why the fecundity-longevity relationship is positive in other systems. In eusocial insects, it has been hypothesised that positive fecundity-longevity relationships could be explained by intrinsically high-quality (i.e. well-resourced) individuals being able to reproduce at higher rates and live longer than low-quality individuals^22,23^. Previous studies in *D. melanogaster* have shown that increased larval diet quality can increase female fertility and longevity^38,39,45^, and the current study has now shown that larval diet quality can change the directionality of the fecundity-longevity relationship. Therefore, asymmetric provisioning, and changes in larval diet quality, could also help explain why eusocial insect reproductives show a positive fecundity-longevity relationship.

Although the results showed that larval diet quality affected the directionality of the fecundity-longevity relationship in adult females, the specific patterns observed differed from those predicted, i.e., that the fecundity-longevity relationship would be negative in L and M females and less negative or positive in H females. One possibility is that the females reared on the high-quality diet had a greater early reproductive investment and a lower mortality rate (as the H females had significantly higher early-life egg production than the M females (Figure 3a) and lower mortality early in the experiment, albeit non-significantly so). This could mean that, within the H treatment, even poorly performing individuals (in terms of longevity) were still able to live beyond the first few weeks because of the higher quality diet. However, this explanation assumes that even low-quality individuals can increase their reproduction without costs on high-quality diets, which would need to be tested.

Another possible explanation comes from the potential cost to individuals of a mismatch between the larval and adult diets. For example, *D. melanogaster* females transferred from a low-quality (20% SYA) larval diet to a high-quality (120% SYA) adult diet exhibited improved fecundity and longevity compared to females transferred from a high-quality larval diet to a low-quality adult diet^45^. However, such an effect seems unlikely to account for the results in the current study because: 1) the sizes of the diet changes for the treatments showing the largest effect size (H and M) were modest (i.e., H females went from a 120% SYA larval diet to a 110% SYA adult diet and M females went from a 100% SYA larval diet to a 110% SYA adult diet); and 2) the treatment females with the largest dietary mismatch (L treatment: 20% SYA larval diet to 110% SYA adult diet) had a flat fecundity-longevity relationship rather than a relationship opposite to those of any of the other two treatments.

A third explanation is that the females used in the current experiment experienced the M diet (i.e., 100% SYA) as the highest quality diet. This is conceivable given that the M diet is the standard diet for *D. melanogaster* rearing^47,48^, the Dahomey populations used to produce flies for this study have been reared on the M diet for more than 20 years (>480 generations), and *D. melanogaster* show evolved responses to their rearing diet^69^. This could explain why females on the M diet had a positive or flat fecundity-longevity relationship, whereas females on the H diet had a negative fecundity-longevity relationship.

An unexpected finding was the non-linear relationship between diet quality and the slope of the fecundity-longevity relationship (i.e., L: non-significant; M: significantly positive or non-significant; H: significantly negative). Previous studies in *D. melanogaster* reported mixed and sometimes inconsistent effects of diet on fecundity and longevity across studies (see Introduction), and complex non-linear effects of diet on fecundity and longevity within studies^39,47,70^. Low-quality diets may reduce fecundity and longevity due to starvation but can also improve longevity due to the longevity-enhancing dietary-restriction effect^33–40^. High-quality diets can provide additional resources for individuals to invest, and therefore increase fecundity and longevity, but high-quality diets can also reduce fecundity and longevity due to toxicity effects^37,40,47^. Our results suggest that these complex and non-linear interacting effects of diet may also affect the directionality of the fecundity-longevity relationship in a non-linear manner.

### Effect of larval diet treatments on gene expression in adult females

The gene expression analyses showed that, in addition to differing in the directionality of the fecundity-longevity relationship, M and H treatments differed markedly from one another in exhibiting: 1) different gene expression profiles across all three sequenced tissues; 2) dissimilar GO terms; and 3) marked contrasts in their degree of overlap with sets of ageing-related genes previously reported in *D. melanogaster*. Interestingly, the strongest gene expression differences between treatments occurred in head and fat body (head: 41.4% up-regulated DEGs shared; 37.4% down-regulated DEGs shared; fat body: 35% up-regulated DEGs shared; 35.7% down-regulated DEGs shared), whereas gene expression was more similar between treatments in the ovaries (80.4% up-regulated DEGs shared; 77.5% down-regulated DEGs shared). These data suggest that the differences between treatments in the fecundity-longevity relationship could be related to large-scale age-related changes in gene expression in head and fat body, and smaller-scale changes in the ovaries. The biological functions of the DEGs also showed pronounced differences between each tissue. For example, in head, in the M treatment, down-regulated DEGs were enriched for mitochondrial processes GO terms, whereas in the H treatment down-regulated DEGs were enriched for transport and nervous system GO terms. In the fat body, in the M treatment, up-regulated DEGs were enriched for immune responses, whereas in the H treatment, up-regulated DEGs were enriched for regulation of the cell cycle.

Across both M and H treatments, we found numerous significant overlaps of age-related DEGs with relevant gene lists from previous studies, namely: 1) age-related DEGs in two previous studies of *D. melanogaster*^66,67^; 2) ageing-related genes in *D. melanogaster* from the GenAge dataset^68^; and 3) the TI-J-LiFe network^32^. These comparisons show inter- and intra-specific commonalities in the genes that change expression with age (and that potentially affect ageing). However, the significant overlaps were markedly different for each treatment. For example, comparison with age-related genes in previous studies of *D. melanogaster* showed significant overlaps in 7/8 comparisons (ranging from 10.2 – 21.2% of genes overlapping). However, in head, the overlaps were greater in the H treatment than in the M treatment (M upregulated = 10.9%, M downregulated = 10.2%; H upregulated = 16%, H down-regulated = 19.4%) whereas in fat body, the overlaps were greater in the M treatment than in the H treatment (M upregulated = 21.2%, M downregulated = 12.7%; H upregulated = no significant overlap, H down-regulated = 11.4%). For the GenAge dataset the largest (and only significant) overlaps with our data were up-regulated genes in the M treatment in head and fat body. For genes in the TI-J-LiFe network, all except one of the comparisons were not significant. However, the one significant comparison involved DEGs in the M treatment, which showed a high overlap of 48% with the TI-J-LiFe gene-list. These comparisons demonstrate that, as might be expected, female *D. melanogaster* exhibit overlaps in their expression of age-related genes across different studies. Further, they also show that larval diet can have a strong effect on age-related gene expression of adults, and, together with our life-history data, indicate that individuals are sensitive to diet quality, and can adjust their gene expression and life-history traits to maintain their fitness on different diets. It remains to be tested whether the observed changes in the directionality of the fecundity-longevity relationship are caused by changes in gene expression, or whether the gene expression changes we observe are responding to the changes in the directionality of the fecundity-longevity relationship.

In conclusion, this study shows that experimental manipulation of larval diet can change the directionality of the fecundity-longevity relationship in adult female *D. melanogaster* and that such changes are accompanied by changes in expression of age- and ageing-related genes. These results were observed even under relatively modest differences in larval diets and even when mean fertility and longevity showed minimal differences between treatments. As well as demonstrating the plasticity of fecundity-longevity relationships, these findings suggest possible mechanistic causes of otherwise puzzling positive fecundity-longevity relationships in other systems, notably eusocial insects. Hence a key outstanding question is whether larval diet quality also affects the plasticity of fecundity-longevity relationships in these other systems.

## Data availability

Data on life history, behaviour, morphometrics, etc., from this study are available in Collins et al.^71^. The raw mRNA-seq sequencing data from this article have been deposited in the National Center for Biotechnology Information’s (NCBI’s) Gene Expression Omnibus (GEO) at https://www.ncbi.nlm.nih.gov/geo/ and are available under series accession number GSE175623. The code used for the analyses are available at https://github.com/davidprince84/Collins-et-al_NE-R000875-1_obj3_mRNA-seq_scripts.

## Acknowledgements

This work was funded by the UK’s Natural Environment Research Council (NERC research grant reference number NE/R000875/1). Laboratory work (library preparation and sequencing) was supported and performed by the NERC Biomolecular Analysis Facility (NBAF) at the University of Edinburgh (Edinburgh Genomics). Edinburgh Genomics is partly supported through core grants from BBSRC (BB/T017864/1) and NERC (UKSBS PR18037). The gene expression analysis presented in this paper was carried out on the High Performance Computing Cluster supported by the Research and Specialist Computing Support service at the University of East Anglia.

## Author Contributions

D.H.C., D.C.P., T.C., and A.F.G.B. designed the study. D.H.C., J.L.D, and D.C.P. performed the experimental manipulations and mRNA-seq sample collections. D.H.C. and D.C.P. analysed the data. D.H.C., D.C.P., and A.F.G.B. wrote the manuscript and all authors commented on drafts. T.C. and A.F.G.B raised funding.

## References

1. Hamilton WD. The moulding of senescence by natural selection. J Theor Biol. 1966;12(1):12–45. doi:10.1016/0022-5193(66)90184-6

2. Flatt T, Partridge L. Horizons in the evolution of aging. BMC Biol. 2018;16(1):1–13. doi:10.1186/s12915-018-0562-z

3. Stearns SC. The Evolution of Life Histories. Oxford University Press, USA; 1992.

4. Flatt T. Survival costs of reproduction in Drosophila. Exp Gerontol. 2011;46(5):369–375. doi:10.1016/j.exger.2010.10.008

5. Maklakov AA, Immler S. The expensive germline and the evolution of ageing. Current Biology. 2016;26(13):R577–R586. doi:10.1016/j.cub.2016.04.012

6. Williams GC. Natural selection, the costs of reproduction, and a refinement of Lack’s principle. Am Nat. 1966;100(916):687–690. doi:10.1086/282461

7. Maklakov AA, Chapman T. Evolution of ageing as a tangle of trade-offs: energy versus function. Proc R Soc B: Biol Sci. 2019;286(1911). doi:10.1098/rspb.2019.1604

8. Clancy DJ, Gems D, Harshman LG, et al. Extension of life-span by loss of CHICO, a Drosophila insulin receptor substrate protein. Science (1979). 2001;292(5514):104–106. doi:10.1126/science.1057991

9. Tu MP, Epstein D, Tatar M. The demography of slow aging in male and female Drosophila mutant for the insulin-receptor substrate homologue chico. Aging Cell.2002;1(1):75–80. doi:10.1046/j.1474-9728.2002.00010.x

10. Marden JH, Rogina B, Montooth KL, Helfand SL. Conditional tradeoffs between aging and organismal performance of Indy long-lived mutant flies. Proc Natl Acad Sci USA. 2003;100(6):3369–3373. doi:10.1073/pnas.0634985100

11. Simon AF, Shih C, Mack A, Benzer S. Steroid control of longevity in Drosophila melanogaster. Science (1979). 2003;299(5611):1407–1410. doi:10.1126/science.1080539

12. Hwangbo DS, Garsham B, Tu MP, Palmer M, Tatar M. Drosophila dFOXO controls lifespan and regulates insulin signalling in brain and fat body. Nature. 2004;429(6991):562–566. doi:10.1038/nature02549

13. Korb J, Heinze J. Ageing and sociality: why, when and how does sociality change ageing patterns? Philos Trans R Soc Lond B Biol Sci. 2021;376(1823):20190727. doi:10.1098/rstb.2019.0727

14. van Noordwijk AJ, de Jong G. Acquisition and allocation of resources: their influence on variation in life history tactics. Am Nat. 1986;128(1):137–142.

15. Reznick D, Nunney L, Tessier A. Big houses, big cars, superfleas and the costs of reproduction. Trends Ecol Evol. 2000;15(10):421–425. doi:10.1016/S0169-5347(00)01941-8

16. Hartmann A, Heinze J. Lay eggs, live longer: division of labor and life span in a clonal ant species. Evolution. 2003;57(10):2424–2429. doi:10.1111/j.0014-3820.2003.tb00254.x

17. Lopez-Vaamonde C, Raine N, Koning J, et al. Lifetime reproductive success and longevity of queens in an annual social insect. J Evol Biol. 2009;22(5):983–996. doi:10.1111/j.1420-9101.2009.01706.x

18. Tsuji K, Kikuta N, Kikuchi T. Determination of the cost of worker reproduction via diminished life span in the ant Diacamma sp. Evolution. 2012;66(5):1322–1331. doi:10.1111/j.1558-5646.2011.01522.x

19. Heinze J, Frohschammer S, Bernadou A. Queen life-span and total reproductive success are positively associated in the ant Cardiocondyla cf. kagutsuchi. Behav Ecol Sociobiol. 2013;67(10):1555–1562. doi:10.1007/s00265-013-1567-9

20. Dixon L, Kuster R, Rueppell O. Reproduction, social behavior, and aging trajectories in honeybee workers. Age. 2014;36(1):89–101. doi:10.1007/s11357-013-9546-7

21. Rueppell O, Königseder F, Heinze J, Schrempf A. Intrinsic survival advantage of social insect queens depends on reproductive activation. J Evol Biol. 2015;28(12):2349–2354. doi:10.1111/jeb.12749

22. Blacher P, Huggins TJ, Bourke AF. Evolution of ageing, costs of reproduction and the fecundity–longevity trade-off in eusocial insects. Proc R Soc B: Biol Sci.2017;284(1858): 20170380. doi:10.1098/rspb.2017.0380

23. Lucas ER, Keller L. The co-evolution of longevity and social life. Funct Ecol.2020;34(1):76–87. doi:10.1111/1365-2435.13445

24. Keller L, Genoud M. Extraordinary lifespans in ants: a test of evolutionary theories of ageing. Nature. 1997;389(6654):958–960. doi:10.1038/40130

25. Heinze J, Schrempf A. Aging and reproduction in social insects - A mini-review. Gerontology. 2008;54(3):160–167. doi:10.1159/000122472

26. Kuhn JMM, Korb J. Editorial overview: Social insects: aging and the re-shaping of the fecundity/longevity trade-off with sociality. Curr Opin Insect Sci. 2016;16:vii–x. doi:10.1016/j.cois.2016.06.002

27. Heinze J, Schrempf A. Terminal investment: individual reproduction of ant queens increases with age. PLoS One. 2012;7(4):e35201. doi:10.1371/journal.pone.0035201

28. Kramer BH, Schrempf A, Scheuerlein A, Heinze J. Ant colonies do not trade-off reproduction against maintenance. PLoS One. 2015;10(9):e0137969. doi:10.1371/journal.pone.0137969

29. Schrempf A, Giehr J, Röhrl R, Steigleder S, Heinze J. Royal Darwinian demons: enforced changes in reproductive efforts do not affect the life expectancy of ant queens. Am Nat. 2017;189(4):436–442. doi:10.1086/691000

30. Jaimes-Niño LM, Heinze J, Oettler J. Late-life fitness gains and reproductive death in Cardiocondyla obscurior ants. Elife. 2022;11:e74695. doi:10.7554/eLife.74695

31. Yan H, Opachaloemphan C, Carmona-Aldana F, et al. Insulin signaling in the long-lived reproductive caste of ants. Science (1979). 2022;377(6610):1092–1099. doi:10.1126/science.abm8767

32. Korb J, Meusemann K, Aumer D, et al. Comparative transcriptomic analysis of the mechanisms underpinning ageing and longevity in social insects. Philos Trans R Soc LondB Biol Sci. 2021;376:20190728. doi:10.1098/rstb.2019.0728

33. Chapman T, Partridge L. Female fitness in Drosophila melanogaster: An interaction between the effect of nutrition and of encounter rate with males. Proc R Soc B: Biol Sci. 1996;263(1371):755–759. doi:10.1098/rspb.1996.0113

34. Partridge L, Piper MDW, Mair W. Dietary restriction in Drosophila. Mech Ageing Dev. 2005;126(9 SPEC. ISS.):938–950. doi:10.1016/j.mad.2005.03.023

35. Metaxakis A, Partridge L. Dietary Restriction Extends Lifespan in Wild-Derived Populations of Drosophila melanogaster. PLoS One. 2013;8(9). doi:10.1371/journal.pone.0074681

36. Semaniuk U, Feden’ko K, Yurkevych IS, Storey KB, Simpson SJ, Lushchak O. Within-diet variation in rates of macronutrient consumption and reproduction does not accompany changes in lifespan in Drosophila melanogaster. Entomol Exp Appl.2018;166(1):74–80. doi:10.1111/eea.12643

37. Lee KP, Simpson SJ, Clissold FJ, et al. Lifespan and reproduction in Drosophila: New insights from nutritional geometry. Proc Natl Acad Sci USA. 2008;105(7):2498–2503. doi:10.1073/pnas.0710787105

38. May CM, Doroszuk A, Zwaan BJ. The effect of developmental nutrition on life span and fecundity depends on the adult reproductive environment in Drosophila melanogaster. Ecol Evol. 2015;5(6):1156–1168. doi:10.1002/ece3.1389

39. May CM, Zwaan BJ. Relating past and present diet to phenotypic and transcriptomic variation in the fruit fly. BMC Genomics. 2017;18(1):1–17. doi:10.1186/s12864-017-3968-z

40. Stefana MI, Driscoll PC, Obata F, et al. Developmental diet regulates Drosophila lifespan via lipid autotoxins. Nat Commun. 2017;8(1). doi:10.1038/s41467-017-01740-9

41. Zandveld J, van den Heuvel J, Mulder M, et al. Pervasive gene expression responses to a fluctuating diet in Drosophila melanogaster: The importance of measuring multiple traits to decouple potential mediators of life span and reproduction. Evolution.2017;71(11):2572–2583. doi:10.1111/evo.13327

42. Pletcher SD, Macdonald SJ, Marguerie R, et al. Genome-wide transcript profiles in aging and calorically restricted Drosophila melanogaster. Current Biology. 2002; 12(9):712–723. doi:10.1016/S0960-9822(02)00808-4

43. Doroszuk A, Jonker MJ, Pul N, Breit TM, Zwaan BJ. Transcriptome analysis of a long-lived natural Drosophila variant: a prominent role of stress-and reproduction genes in lifespan extension. BMC Genomics. 2012;13(1). doi:10.1186/1471-2164-13-167

44. Partridge L, Alic N, Bjedov I, Piper MDW. Ageing in Drosophila: The role of the insulin/Igf and TOR signalling network. Exp Gerontol. 2011;46(5):376–381. doi:10.1016/j.exger.2010.09.003

45. Duxbury EML, Chapman T. Sex-Specific Responses of Life Span and Fitness to Variation in Developmental Versus Adult Diets in Drosophila melanogaster. J Gerontol A Biol Sci Med Sci. 2020;75(8):1431–1438. doi:10.1093/gerona/glz175

46. Tu MP, Tatar M. Juvenile diet restriction and the aging and reproduction of adult Drosophila melanogaster. Aging Cell. 2003;2(6):327–333. doi:10.1046/j.1474-9728.2003.00064.x

47. Bass TM, Grandison RC, Wong R, Martinez P, Partridge L, Piper MDW. Optimization of Dietary Restriction Protocols in Drosophila. J Gerontol A Biol Sci Med Sci.2007;62(10):1071–1081.

48. Magwere T, Chapman T, Partridge L. Sex Differences in the Effect of Dietary Restriction on Life Span and Mortality Rates in Female and Male Drosophila Melanogaster. J Gerontol A Biol Sci Med Sci. 2004;59(1):3–9. doi:10.1093/gerona/59.1.b3

49. Shukla AK, Johnson K, Giniger E. Common features of aging fail to occur in Drosophila raised without a bacterial microbiome. iScience. 2021;24(7):102703. doi:10.1016/j.isci.2021.102703

50. R core team. A language and environment for statistical computing. R Foundation for Statistical Computing, Vienna, Austria. URL https://www.R-project.org/. Published online 2020.

51. Bates D, Mächler M, Bolker BM, Walker SC. Fitting linear mixed-effects models using lme4. J Stat Softw. 2015;67(1). doi:10.18637/jss.v067.i01

52. Brooks ME, Kristensen K, van Benthem KJ, et al. glmmTMB balances speed and flexibility among packages for zero-inflated generalized linear mixed modeling. R Journal. 2017;9(2):378–400. doi:10.32614/rj-2017-066

53. Therneau TM. A Package for Survival Analysis in R. R package version 3.4-0, https://CRAN.R-project.org/package=survival.

54. Kleiber C, Zeileis A. Applied Econometrics with R. Springer-Verlag; 2008.

55. Andrews S. FastQC: a quality control tool for high throughput sequence data. Available online at: http://www.bioinformatics.babraham.ac.uk/projects/fastqc.

56. Ewels P, Magnusson M, Lundin S, Käller M. MultiQC: Summarize analysis results for multiple tools and samples in a single report. Bioinformatics. 2016;32(19):3047–3048. doi:10.1093/bioinformatics/btw354

57. van Rossum G, Drake F. Python 3 Reference Manual. CreateSpace; 2009.

58. Kim D, Langmead B, Salzberg S. HISAT: a fast spliced aligner with low memory requirements. Nat Methods. 2015;12:357–360. doi:10.1038/nmeth.3317

59. Wang L, Wang S, Li W. RSeQC: Quality control of RNA-seq experiments. Bioinformatics. 2012;28(16):2184–2185. doi:10.1093/bioinformatics/bts356

60. Bray NL, Pimentel H, Melsted P, Pachter L. Near-optimal probabilistic RNA-seq quantification. Nat Biotechnol. 2016;34(5):525–527. doi:10.1038/nbt.3519

61. Love MI, Huber W, Anders S. Moderated estimation of fold change and dispersion for RNA-seq data with DESeq2. Genome Biol. 2014;15(12):1–21. doi:10.1186/s13059-014-0550-8

62. Yu G, Wang LG, Han Y, He QY. ClusterProfiler: An R package for comparing biological themes among gene clusters. OMICS. 2012;16(5):284–287. doi:10.1089/omi.2011.0118

63. Carlson M. .org.Dm.eg.db: Genome wide annotation for Fly. R package version 3.11.4.

64. Boyle EI, Weng S, Gollub J, et al. GO::TermFinder - Open source software for accessing Gene Ontology information and finding significantly enriched Gene Ontology terms associated with a list of genes. Bioinformatics. 2004;20(18):3710–3715. doi:10.1093/bioinformatics/bth456

65. Yu G. Gene Ontology Semantic Similarity Analysis Using GOSemSim. In: Kidder B, ed. Methods in Molecular Biology. Humana, New York, NY; 2020:207–215. doi:10.1007/978-1-0716-0301-7_11.

66. Chen H, Zheng X, Zheng Y. Age-associated loss of lamin-B leads to systemic inflammation and gut hyperplasia. Cell. 2014;159(4):829–843. doi:10.1016/j.cell.2014.10.028.

67. Pacifico R, MacMullen CM, Walkinshaw E, Zhang X, Davis RL. Brain transcriptome changes in the aging Drosophila melanogaster accompany olfactory memory performance deficits. PLoS One. 2018;13(12):e0209405. doi: 10.1371/journal.pone.0209405

68. Tacutu R, Thornton D, Johnson E, et al. Human ageing genomic resources: new and updated databases. Nucleic Acids Res. 2018;46(D1):D1083–D1090. doi:10.1093/nar/gkx1042

69. Zajitschek F, Georgolopoulos G, Vourlou A, et al. Evolution under dietary restriction decouples survival from fecundity in Drosophila melanogaster females. J Gerontol A Biol Sci Med Sci. 2019;74(10):1542–1548. doi:10.1093/gerona/gly070

70. Fricke C, Bretman A, Chapman T. Adult male nutrition and reproductive success in Drosophila melanogaster. Evolution. 2008;62(12):3170–3177. doi:10.1111/j.1558-5646.2008.00515.x

71. Collins DH, Prince DC, Donelan JL, Chapman T, Bourke AF. Life history, developmental and morphometric data for individual flies from an experiment manipulating larval nutrition in female fruit flies (Drosophila melanogaster). NERC EDS Environmental Information Data Centre. Published online 2021. doi:https://doi.org/10.5285/90dee4ce-187c-4094-8895-b079d29922f5

