## Supplemental information for "Developmental diet alters the fecundity-longevity relationship and age-related gene expression in *Drosophila melanogaster*"

### Supplemental methods

Effect of treatment on development duration, development success, and adult female thorax size

We estimated the effect of treatment on two measures of development duration, pupation duration (the mean interval in hours per vial between the time at which larvae of either sex were first placed in vials and the start of pupation) and eclosion duration (the mean interval in hours per vial between the time at which larvae of either sex were first placed in vials and the start of eclosion). We used a glm with gamma error distribution and a log link function to determine the effect of each treatment on pupation duration and eclosion duration. For each response variable, the full model contained an additional fixed effect for population (one of the four Dahomey source populations from which larvae in each vial were taken), and each model was compared with and without this term using the second order Akaike Information Criterion (AICc) to determine the best model (Pinheiro & Bates, 2002). In each case, the final model did not include additional fixed effects (adding population changed the AICc value by less than 2 and did not affect the significance of the treatment fixed effect in the model).

We estimated the effect of treatment on pupation success (number of larvae, of both sexes, that pupated out of 100 first-instar larvae that were added to each vial) and eclosion success (number of larvae, of both sexes, that eclosed out of 100 first-instar larvae that were added to each vial) using generalised linear mixed effects models (glmm) with Poisson error distribution and a log link function. Graphical and analytical tests showed that the data were not overdispersed (overdispersion test: *c* = -0.041, *p* = 0.638). For each response variable, the full model contained additional fixed effects for researcher (one of the four researchers who counted out first-instar larvae into each vial) and population (one of the four Dahomey source populations from which larvae in each vial were taken), and each model was compared with and without these terms using AICc. In each case the model with the lowest AICc value included researcher but not population as an additional effect. Therefore the final model included researcher but not population as an additional fixed effect. The final model for each response variable was compared with an intercept-only model with the fixed effects removed using AICc and a likelihood ratio test (LRT) to determine whether the models containing the fixed effects provided the best fit to the data.

Due to the bodies of some females deteriorating after death, we were able to measure thorax size (distance (mm) from the anterior margin of the thorax to the posterior tip of the scutellum, as laterally viewed) for only a sub-sample of adult females from each treatment. Therefore, sample sizes for females with measurements of thorax size were as follows: L (n = 48), M (n = 38), and H (n = 46). To test whether thorax size should be included as a possible predictor effect in the models that investigated the effect of treatment on fertility and longevity, we used a glm (with gaussian error distribution and a log link function) to test whether treatment had a significant effect on thorax size.

### Supplemental results

Effect of treatment on development duration, development success, and adult female thorax size

Treatment significantly affected pupation and eclosion duration: the models that included treatment as a fixed effect had a significantly better fit to the data than the intercept-only models (pupation time: deltaAICc = 363; LRT: χ^2^ = 6.824, df = 2, p < 0.001; eclosion time: deltaAICc = 314.2; LRT: χ^2^ = 2.541, df = 2, p < 0.001). Larvae reared on the L treatment had significantly longer pupation durations (mean across all L (N = 24) vials (SD) = 205 (6.4) hours) than larvae reared on the M treatment (mean across all M (N = 25) vials (SD) = 110 (2.9) hours; gamma glmm: t = 79.463, p < 0.001), and larvae on the M treatment had significantly longer pupation durations than larvae reared on the H treatment (mean across all H (N = 24) vials (SD) = 108 (1.9) hours; gamma glmm: t = 2.436, p = 0.009). In addition, larvae reared on the L treatment had significantly longer eclosion durations (mean across all L vials (SD) = 311 (8.14) hours) than larvae reared on the M treatment (mean across all M vials (SD) = 213 (5.16) hours; glmm: t = 64.176, p < 0.001), and larvae on the M treatment had significantly longer eclosion times than larvae reared on the H treatment (mean across all H vials (SD) = 209 (2.52) hours; glmm: t = 2.938, p = 0.009; Figures S2 – S3).

There was no significant effect of treatment on pupation success or eclosion success as in each case the best model that contained treatment as a fixed effect had a lower AICc value than the intercept-only model and a likelihood ratio test showed no significant difference from the intercept-only model (pupation success: deltaAICc = 1.9; LRT: χ^2^ = 0.408, df = 1, p = 0.523; eclosion success: deltaAICc = 1.8; LRT: χ^2^ = 0.528, df = 1, p = 0.467; Figures S4 – S5).

Treatment had a significant effect on adult female thorax size (gaussian glm: LRT: *χ^2^* = 0.171, df = 2, p < 0.001; Figure S6). L females (N = 48; mean (SD) size = 0.96 (0.043) mm) had significantly smaller thorax size than M females (N = 38; mean (SD) size = 1.04 (0.029) mm; gaussian glm: t = -6.101, p < 0.001) and H females (N = 46; mean (SD) size = 1.04 (0.027) mm; gaussian glm: t = -7.406, p < 0.001); however, M females were not significantly different in thorax size from H females (gaussian glm: t = 0.704, p = 0.484).

### Supplemental Figures

#### Figure_S1

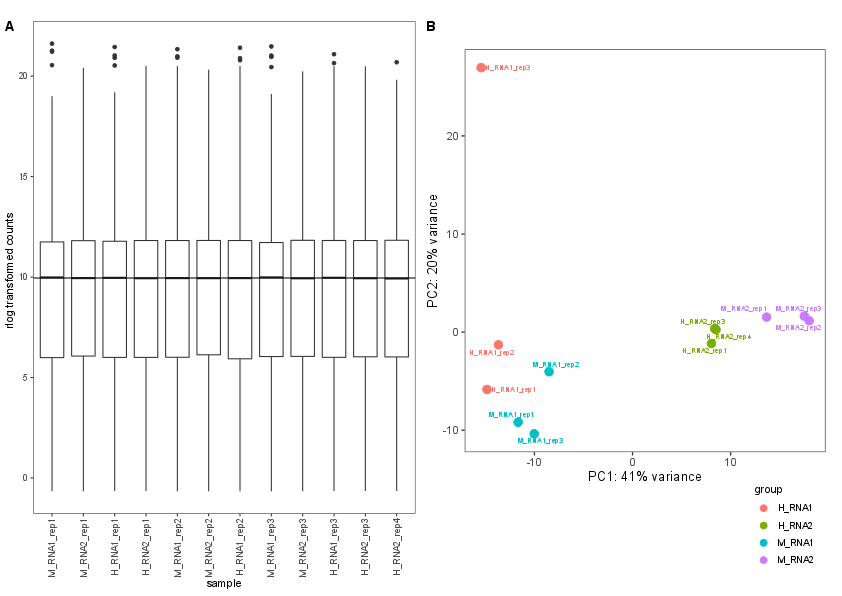

Figure S1. Exploratory plots from the differential expression analysis of mRNA-seq libraries from pooled heads of adult female *Drosophila melanogaster*. (A) Normalisation boxplots of the rlog-transformed value of mRNA-seq expression for genes in each library. Black horizontal bars: medians; boxes: interquartile ranges; whiskers: 10^th^ to 90^th^ percentile ranges. (B) Principal component analysis (PCA) plots for all expressed genes isolated from mRNA-seq libraries in head. Axes represent principal components. Individual points represent biological replicates (coloured by group). Sample names in the format: treatment_time point_biological replicate. M, 100% SYA treatment diet; H, 120% SYA treatment diet; RNA1, time point 1 (after 10% cohort death); RNA2, time point 2 (after 60% cohort death), rep1, biological replicate 1. Head mRNA-seq libraries: N = 12.

#### Figure_S2

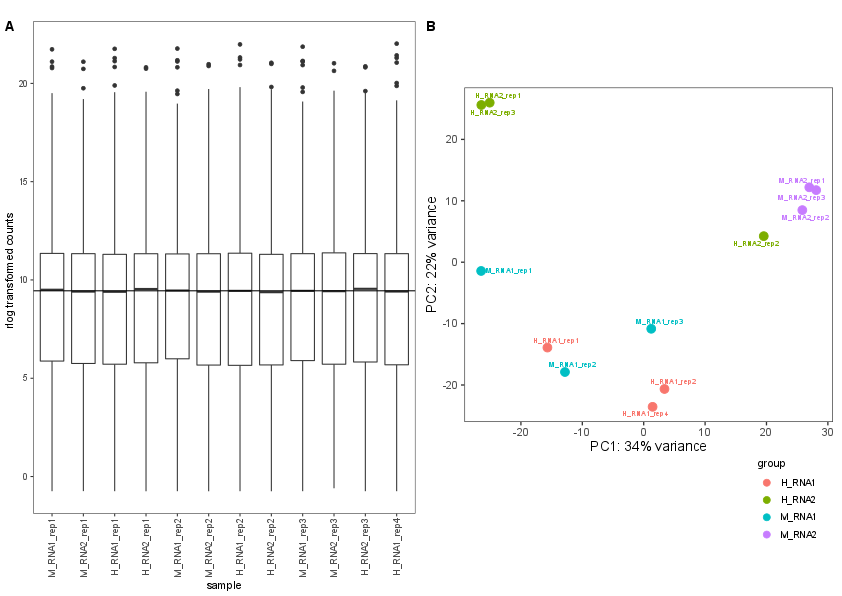

Figure S2. Exploratory plots from the differential expression analysis of mRNA-seq libraries from pooled fat bodies of adult female *Drosophila melanogaster*. (A) Normalisation boxplots of the rlog-transformed value of mRNA-seq expression for genes in each library. Black horizontal bars: medians; boxes: interquartile ranges; whiskers: 10^th^ to 90^th^ percentile ranges. (B) Principal component analysis (PCA) plots for all expressed genes isolated from mRNA-seq libraries in fat body. Axes represent principal components. Individual points represent biological replicates (coloured by group). Sample names in the format: treatment_time point_biological replicate. M, 100% SYA treatment diet; H, 120% SYA treatment diet; RNA1, time point 1 (after 10% cohort death); RNA2, time point 2 (after 60% cohort death), rep1, biological replicate 1. Fat body mRNA-seq libraries: N = 12.

#### Figure_S3

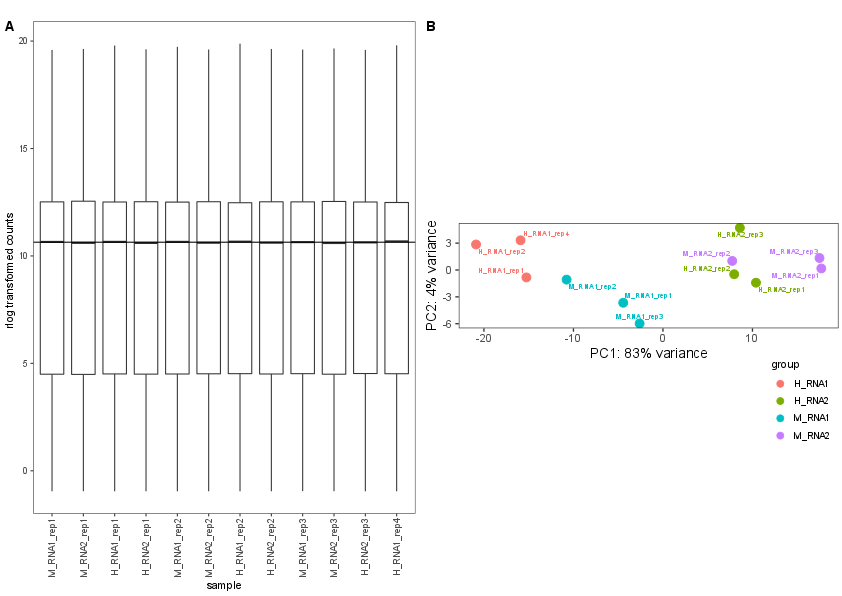

Figure S3. Exploratory plots from the differential expression analysis of mRNA-seq libraries from pooled ovaries of adult female *Drosophila melanogaster*. (A) Normalisation boxplots of the rlog-transformed value of mRNA-seq expression for genes in each library. Black horizontal bars: medians; boxes: interquartile ranges; whiskers: 10^th^ to 90^th^ percentile ranges. (B) Principal component analysis (PCA) plots for all expressed genes isolated from mRNA-seq libraries in ovaries. Axes represent principal components. Individual points represent biological replicates (coloured by group). Sample names in the format: treatment_time point_biological replicate. M, 100% SYA treatment diet; H, 120% SYA treatment diet; RNA1, time point 1 (after 10% cohort death); RNA2, time point 2 (after 60% cohort death), rep1, biological replicate 1. Ovaries mRNA-seq libraries: N = 12.

#### Figure S4

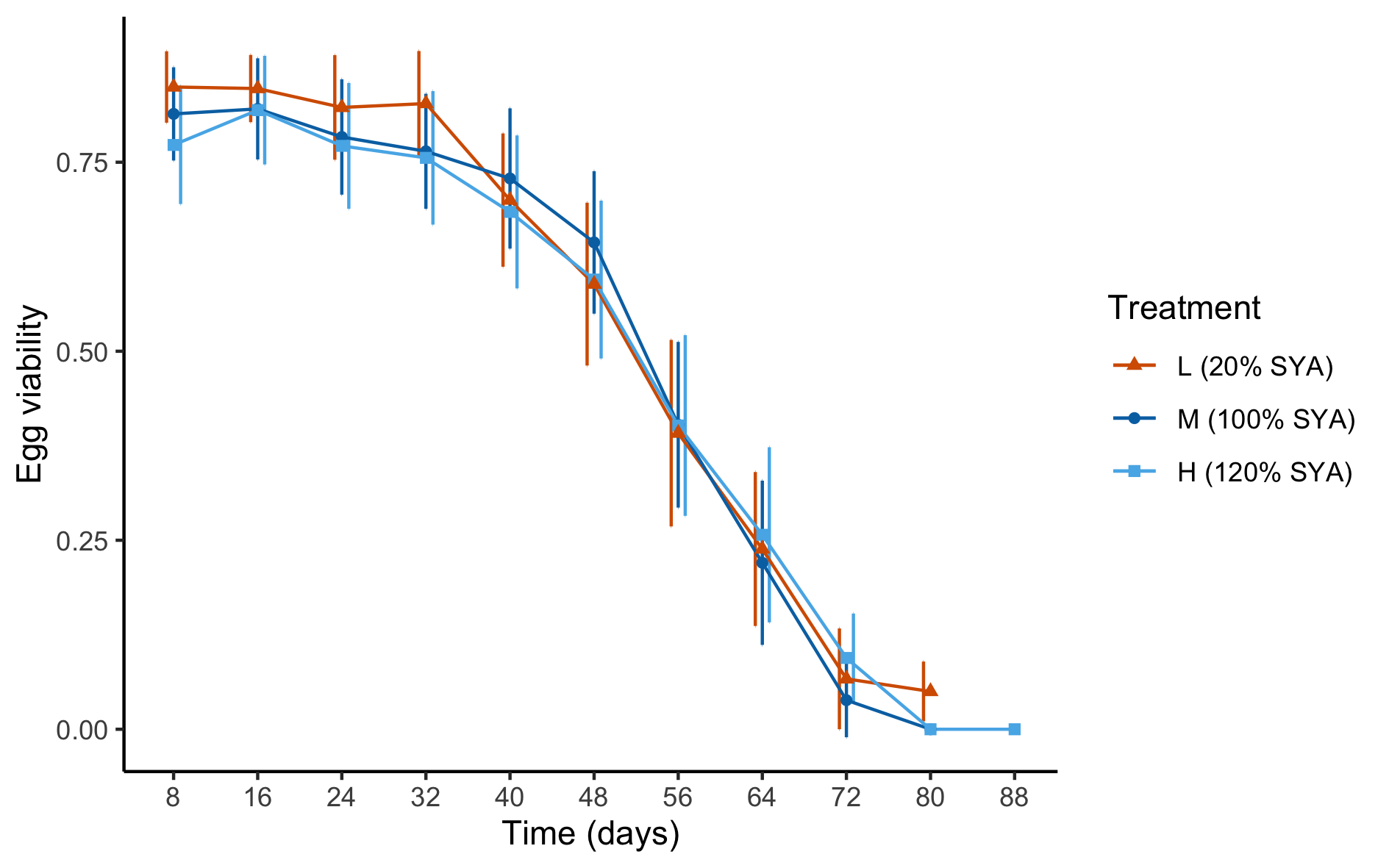

Figure S4. Mean egg viability (whole-life offspring production as a proportion of whole-life egg production) as a function of time (represented by the last day of each successive eight-day cycle) for eggs laid by adult female *Drosophila melanogaster* reared on L (20% SYA, orange triangles, n = 56), M (100% SYA, dark blue circles, n = 45), H (120% SYA, light blue squares, n = 56) treatment diets as larvae then on a 110% SYA diet as adults. Points: mean (per female) egg viability across all females for each experiment (eight-day) cycle; error bars: 1 S.D. Egg viability was not significantly affected by treatment but decreased with time (see Results for details).

#### Figure S5

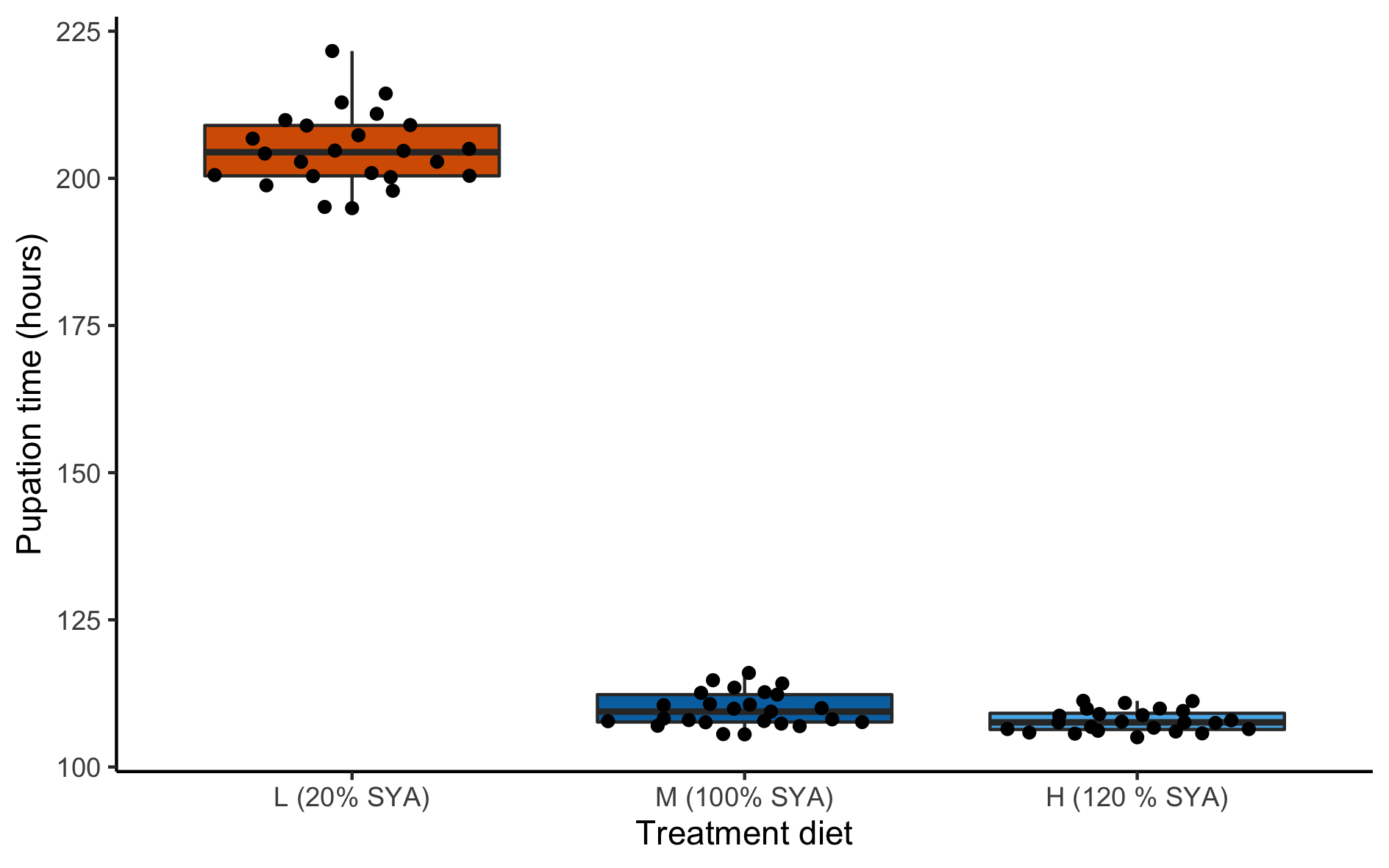

Figure S5. Pupation time (the mean duration in hours per vial that it took larvae of both sexes to pupate) for *Drosophila melanogaster* larvae reared on L (20% SYA, n = 24, orange), M (100% SYA, n = 25, dark blue), and H (120% SYA, n = 24, light blue) diets. Black circles: mean duration for each vial. Horizontal bars: median (across all larvae within each treatment); boxes: interquartile ranges; whiskers: ranges up to 1.5 × the interquartile range. L larvae had significantly longer pupation times than M larvae, and M larvae had significantly longer pupation times than H larvae (see Supplemental results for details).

#### Figure S6

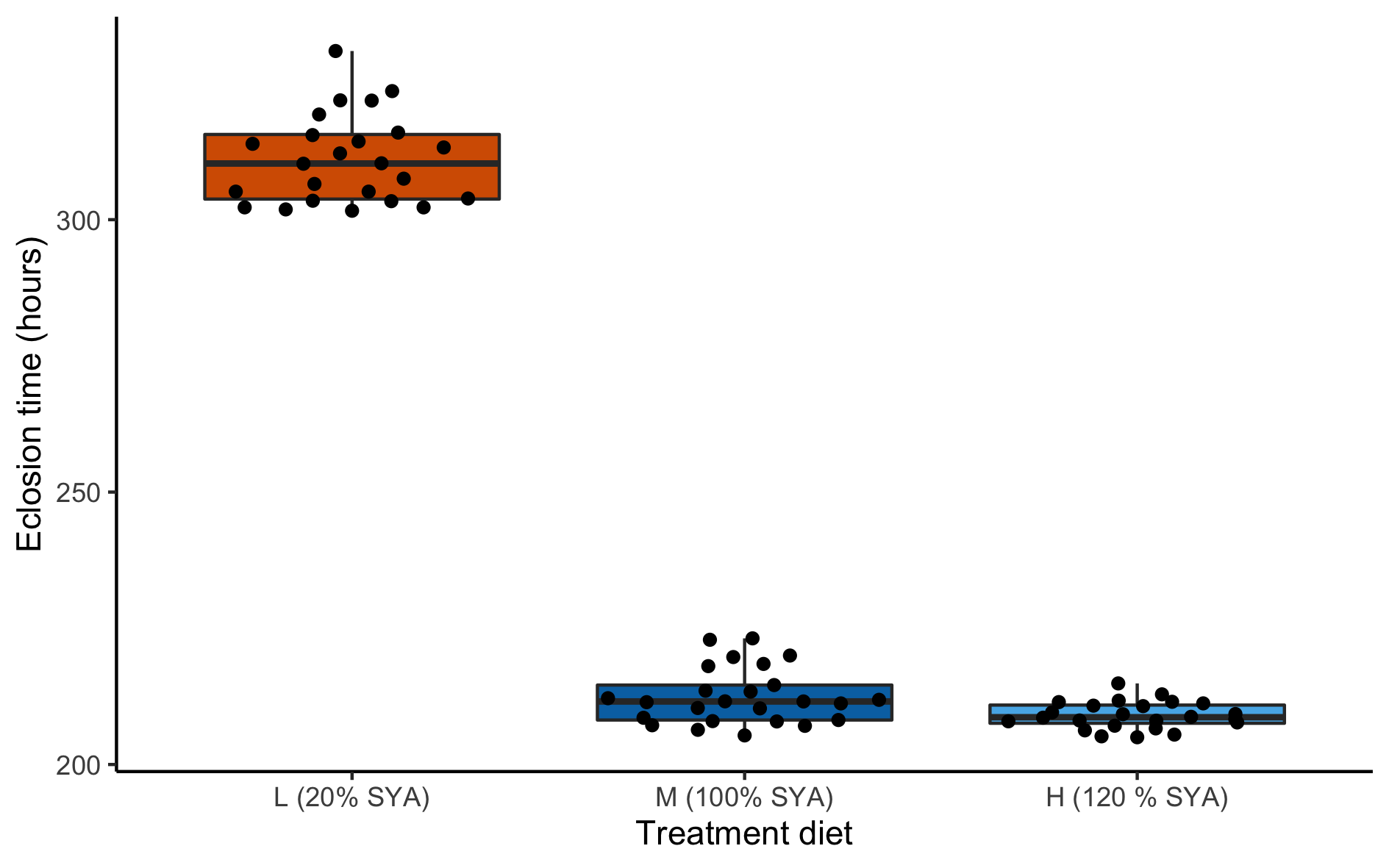

Figure S6. Eclosion time (the mean duration in hours per vial that it took larvae of both sexes to eclose into adults) for *Drosophila melanogaster* larvae reared on L (20% SYA, n = 24, orange), M (100% SYA, n = 25, dark blue), and H (120% SYA, n = 24, light blue) diets. Black circles: mean duration for each vial. Horizontal bars: median (across all larvae within each treatment); boxes: interquartile ranges; whiskers: ranges up to 1.5 × the interquartile range. L larvae had significantly longer eclosion times than M larvae, and M larvae had significantly longer eclosion times than H larvae (see Supplemental results for details).

#### Figure S7

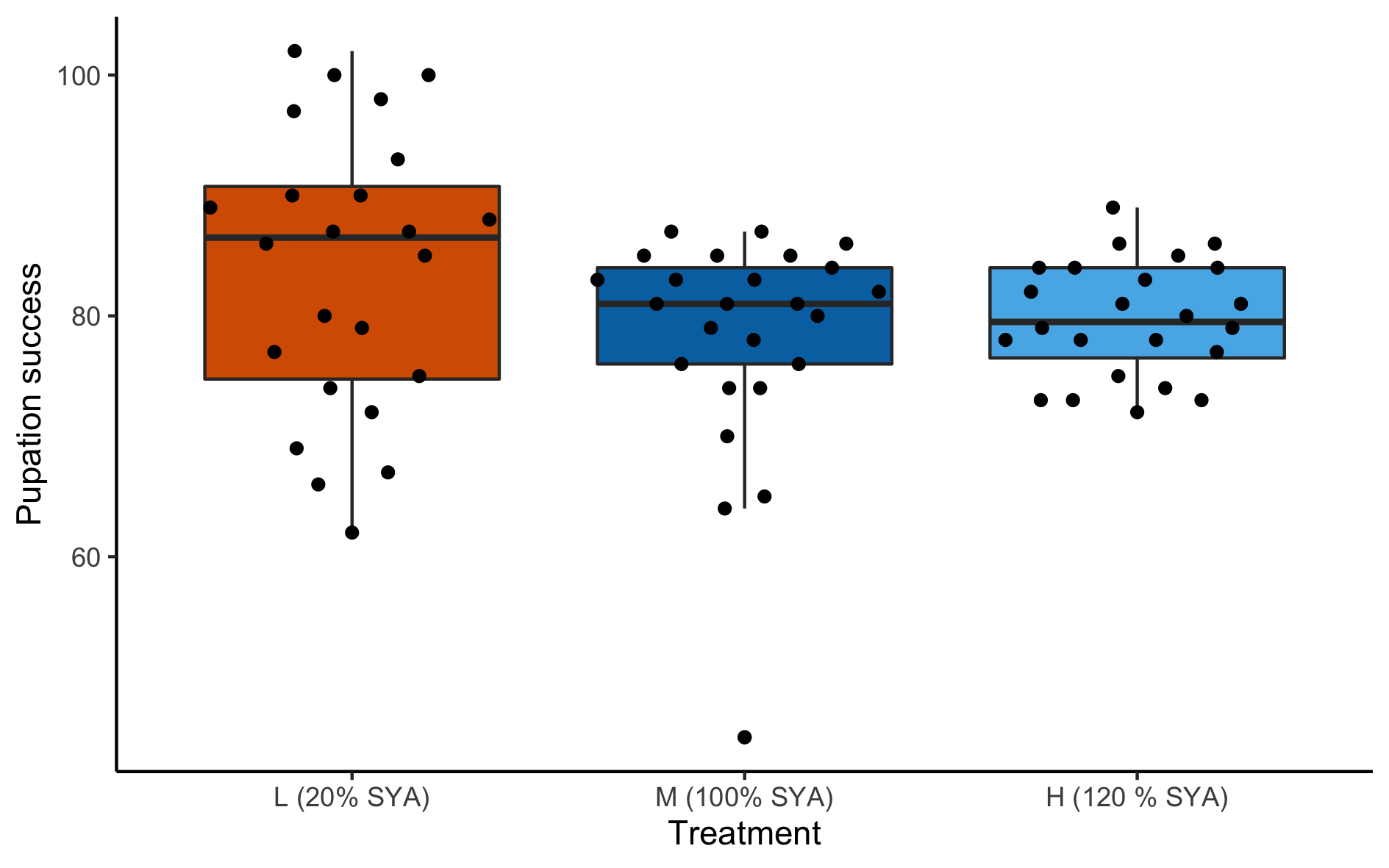

Figure S7. Pupation success (the number of larvae that pupated out of 100 first-instar larvae of both sexes that were added to each vial) for *Drosophila melanogaster* larvae reared on L (20% SYA, n = 24, orange), M (100% SYA, n = 25, dark blue), and H (120% SYA, n = 24, light blue) diets. Black circles: number of pupae in each vial. Horizontal bars: median (across all larvae within each treatment); boxes: interquartile ranges; whiskers: ranges up to 1.5 × the interquartile range. There were no significant differences between treatments in pupation success (see Supplemental results for details).

#### Figure S8

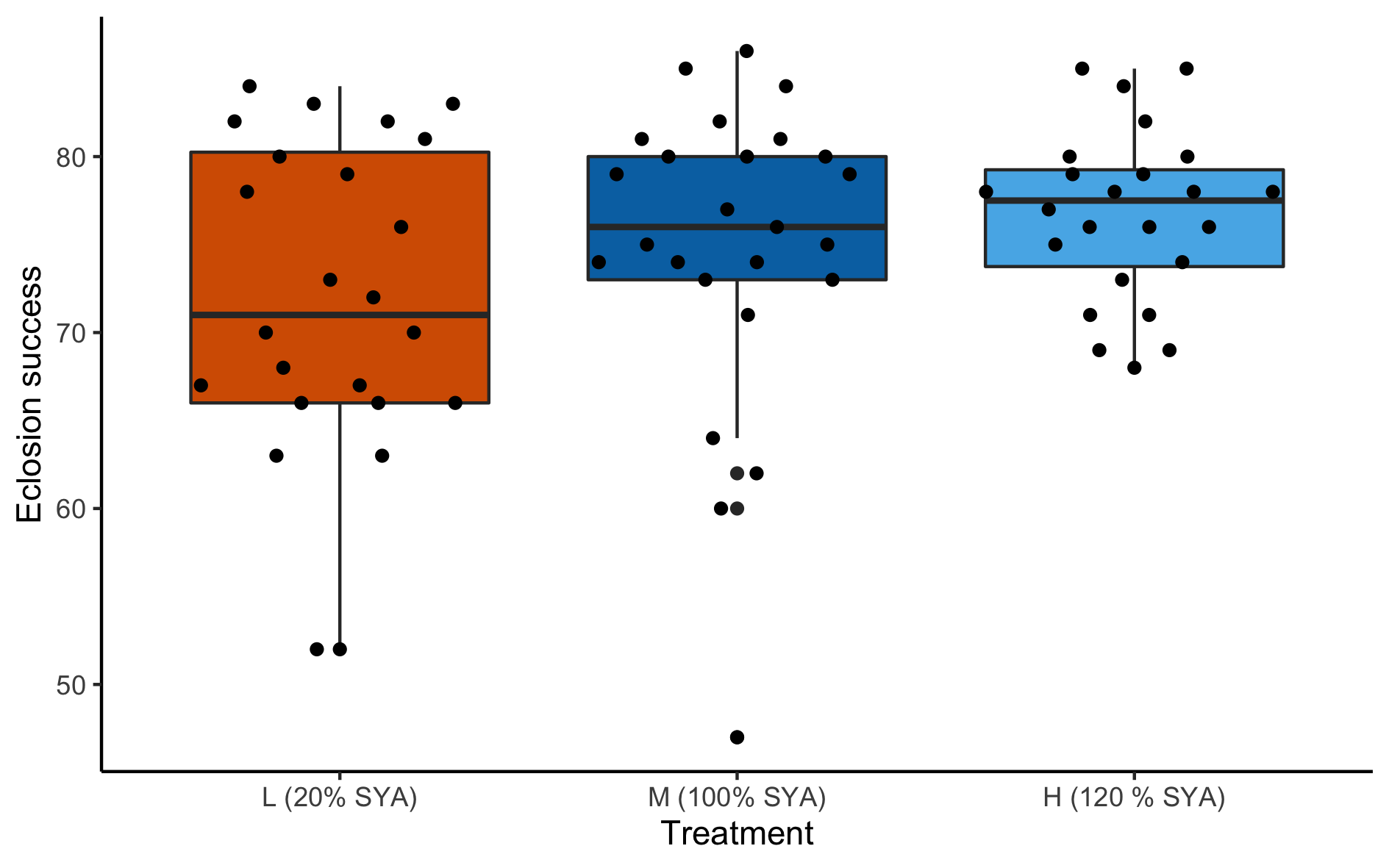

Figure S8. Eclosion success (the number of larvae of both sexes that eclosed into adults out of 100 first-instar larvae that were added to each vial) for *Drosophila melanogaster* larvae reared on L (20% SYA, n = 24, orange), M (100% SYA, n = 25, dark blue), and H (120% SYA, n = 24, light blue) diets. Black circles: number of adults in each vial. Horizontal bars: median (across all larvae within each treatment); boxes: interquartile ranges; whiskers: ranges up to 1.5 × the interquartile range. There were no significant differences between treatments in eclosion success (see Supplemental results for details).

#### Figure S9

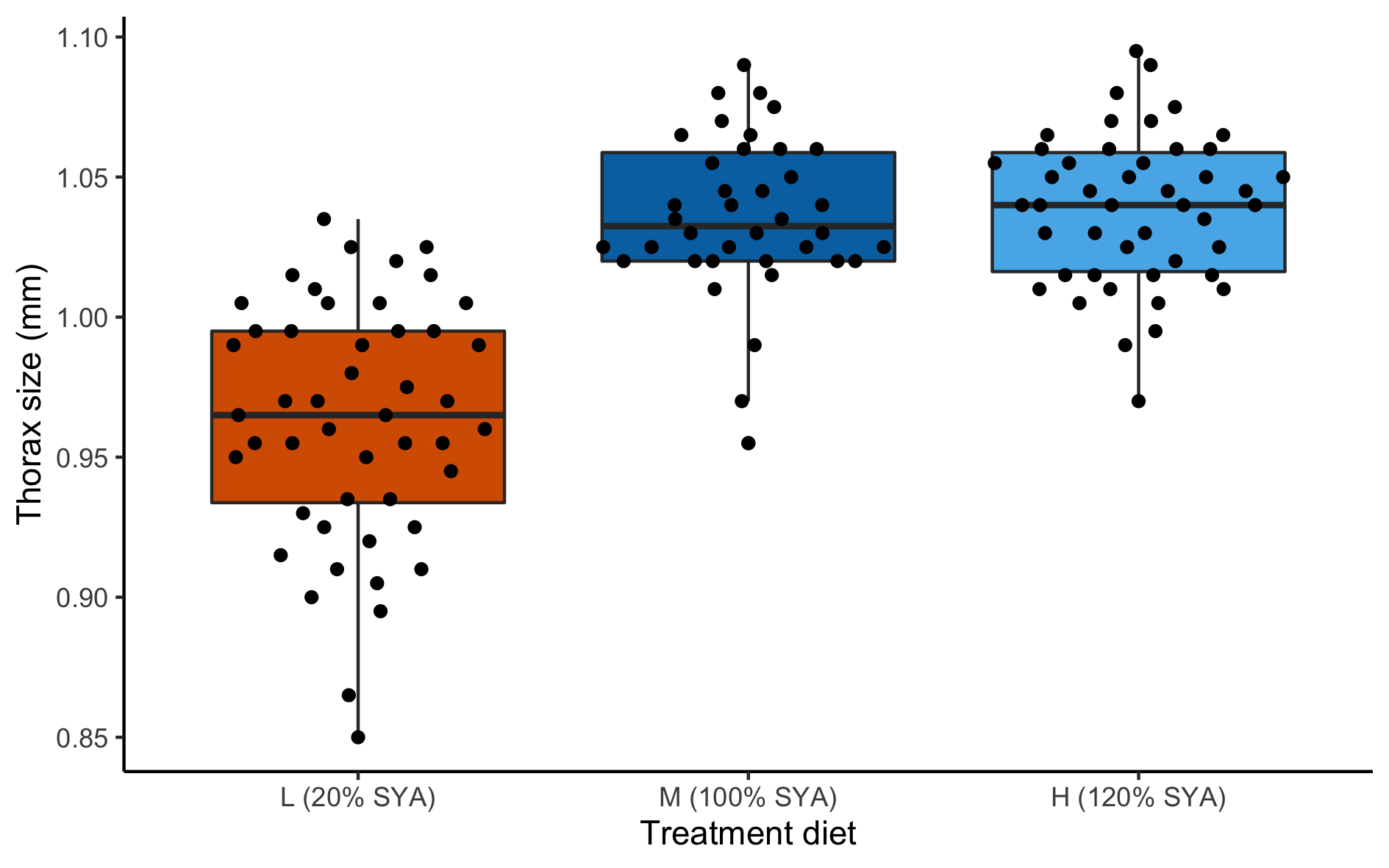

Figure S9. Thorax size (distance (mm) from the anterior margin of the thorax to the posterior tip of the scutellum, as laterally viewed) for adult female *Drosophila melanogaster* reared on L (20% SYA, n = 46, orange), M (100% SYA, n = 38, dark blue), and H (120% SYA, n = 48, light blue) treatment diets as larvae and then on a 110% SYA diet as adults. Horizontal bars: median (across all females within each treatment); boxes: interquartile ranges; whiskers: ranges up to 1.5 × the interquartile range. L females were significantly smaller than M or H females (see Supplemental results for details).

#### Figure S10

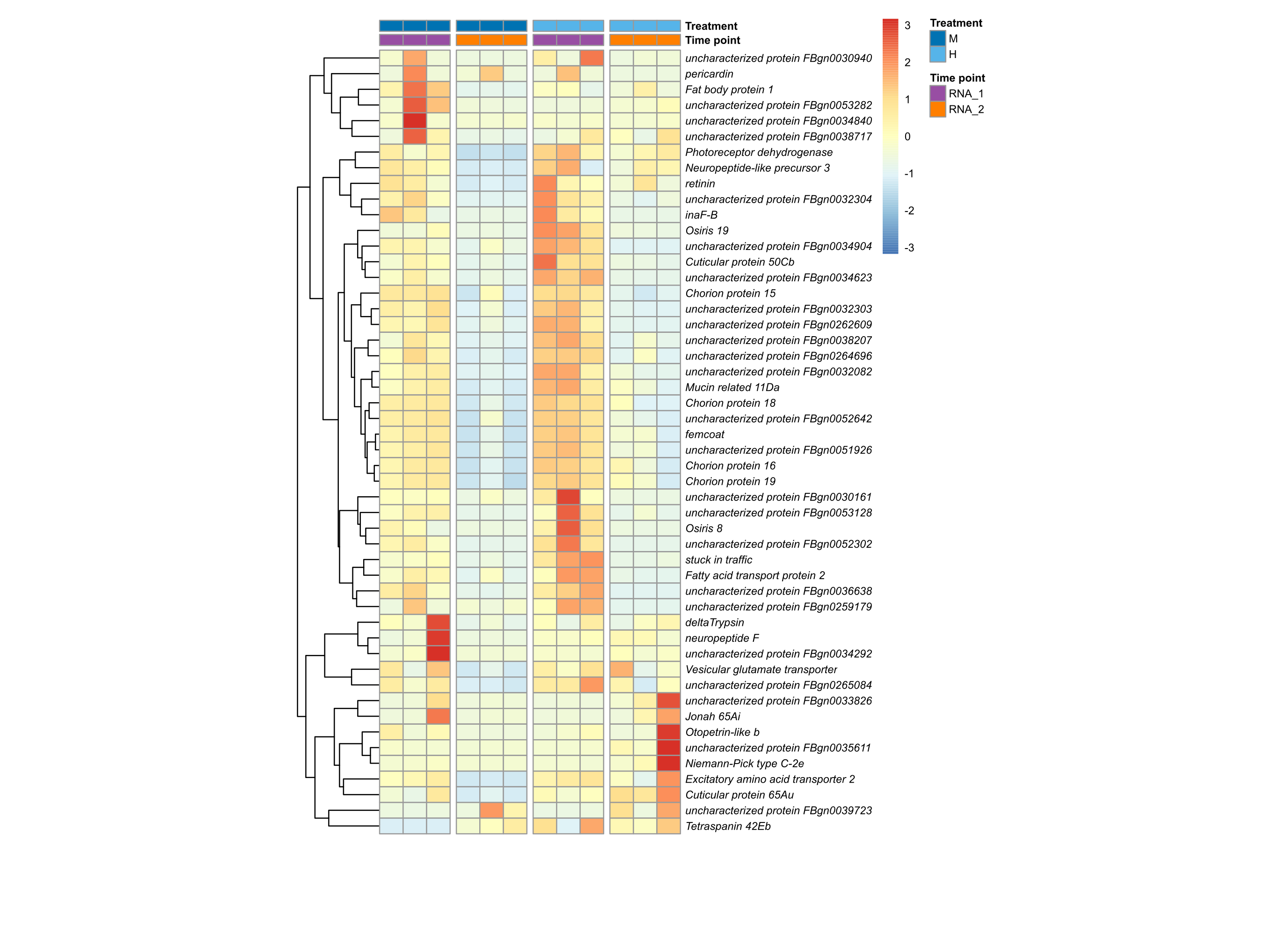

Figure S10. Gene expression differences from mRNA-seq libraries prepared from pooled head samples of adult female *Drosophila melanogaster* from M and H treatments. Differences expressed in M and H treatments at two timepoints (RNA-1, purple and RNA-2, orange) showing relative changes in gene expression (log_2_ fold change) within each gene for the 50 most highly differentially expressed genes (DEGs) (out of 6,797 DEGs in total). Each row represents an individual gene and each column represents a biological replicate of the mRNA-seq data. Vertical breaks represent the two treatments (M (blue) and H (green)) and the two time points (RNA-1 (purple) and RNA-2 (orange)). The dendrogram at left groups genes that cluster according to their gene expression patterns.

#### Figure S11

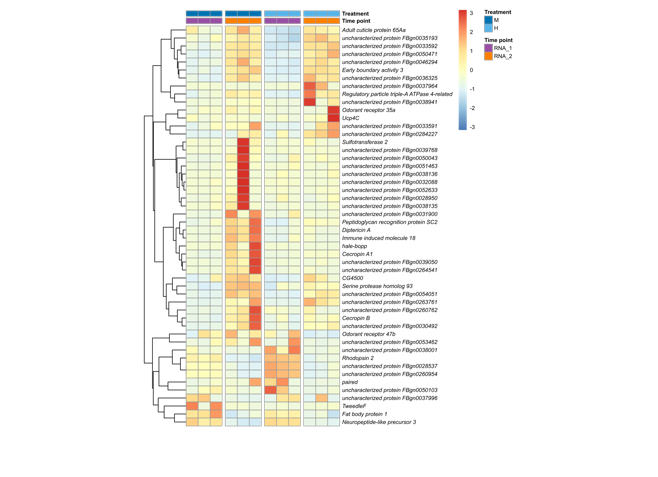

Figure S11. Gene expression differences from mRNA-seq libraries prepared from pooled fat body samples of adult female *Drosophila melanogaster* from M and H treatments. Differences expressed in M and H treatments at two timepoints (RNA-1, purple and RNA-2, orange) showing relative changes in gene expression (log_2_ fold change) within each gene for the 50 most highly differentially expressed genes (DEGs) (out of 5,702 DEGs in total). Each row represents an individual gene and each column represents a biological replicate of the mRNA-seq data. Vertical breaks represent the two treatments (M (blue) and H (green)) and the two time points (RNA-1 (purple) and RNA-2 (orange)). The dendrogram at left groups genes that cluster according to their gene expression patterns.

#### Figure S12

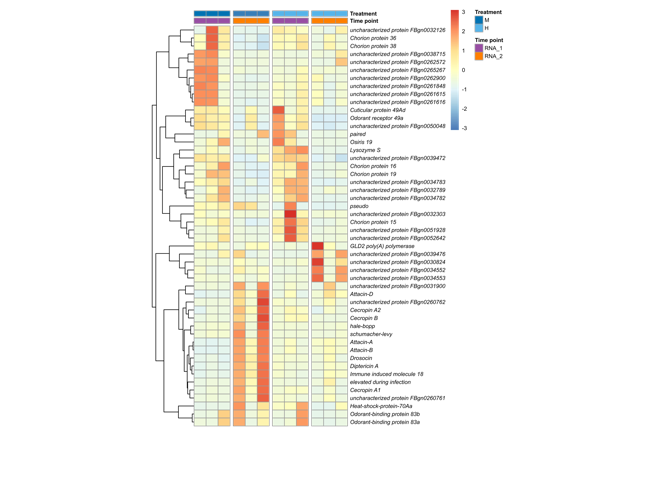

Figure S12. Gene expression differences from mRNA-seq libraries prepared from pooled ovary samples of adult female *Drosophila melanogaster* from M and H treatments. Differences expressed in M and H treatments at two timepoints (RNA-1, purple and RNA-2, orange) showing relative changes in gene expression (log_2_ fold change) within each gene for the 50 most highly differentially expressed genes (DEGs) (out of 7,320 DEGs in total). Each row represents an individual gene and each column represents a biological replicate of the mRNA-seq data. Vertical breaks represent the two treatments (M (blue) and H (green)) and the two time points (RNA-1 (purple) and RNA-2 (orange)). The dendrogram at left groups genes that cluster according to their gene expression patterns.

#### Figure S13

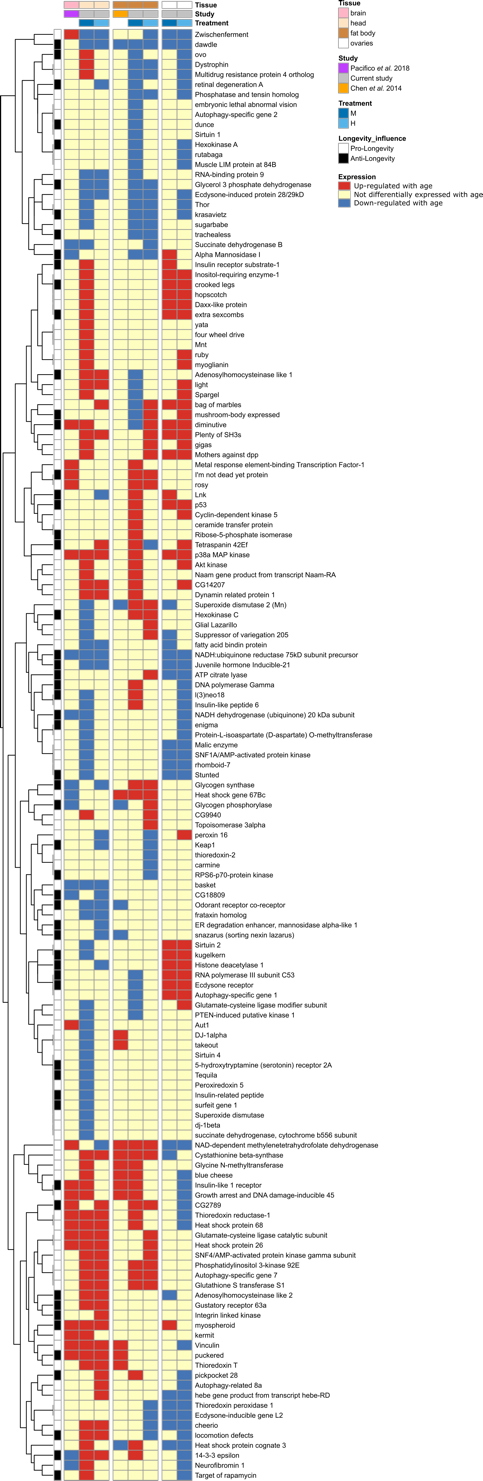

Figure S13. Comparison of age-related genes in three RNAseq datasets of adult female *Drosophila melanogaster* with GenAge genes. The three datasets consist of: 1) three tissues (head, fat body, and ovaries) from M (100% SYA) and H (120% SYA) females in the current study; 2) brain from females in Pacifico et al. (2018); and 3) fat body from females in Chen et al. (2014). Each row represents a gene from *Drosophila melanogaster* in the GenAge database (Tacatu et al. 2018). Boxes next to each gene row represent pro-longevity (white) and anti-longevity (black) genes. Each column shows the age-related expression status of focal genes in a given treatment and tissue in *D. melanogaster* females in the given study. Vertical breaks separate the three tissues studied in this study (head, fat body, and ovaries) and in the two comparison studies (brain, fat body). The dendogram at left groups genes according to their gene expression patterns.
