## Supplemental file S3 for "Developmental diet alters the fecundity-longevity relationship and age-related gene expression in *Drosophila melanogaster*"

MultiQC Report


### Toggle navigation v1.9

Loading report..

- General Stats
- FastQC
  - Sequence Counts
  - Sequence Quality Histograms
  - Per Sequence Quality Scores
  - Per Base Sequence Content
  - Per Sequence GC Content
  - Per Base N Content
  - Sequence Length Distribution
  - Sequence Duplication Levels
  - Overrepresented sequences
  - Adapter Content
  - Status Checks

Toolbox

##### MultiQC Toolbox

###### Apply Highlight Samples

+

Regex mode off
help
 Clear

###### Apply Rename Samples

+

Click here for bulk input.

Paste two columns of a tab-delimited table here (eg. from Excel).

First column should be the old name, second column the new name.

Add

Regex mode off
help
 Clear

###### Apply Show / Hide Samples

Hide matching samples

Show only matching samples

+

Regex mode off
help
 Clear

###### Export Plots

- Images
- Data

px

px

Aspect ratio

PNG
JPEG
SVG

Plot scaling

X

Download the raw data used to create the plots in this report below:

Format:

Tab-separated
Comma-separated
JSON

Note that additional data was saved in `00_NER0008751_obj3_exp3_supplementary_file_S3_ovary_fastqc_multiqc_data` when this report was generated.

---

###### Choose Plots

 All
 None


---


   Download Plot Images

If you use plots from MultiQC in a publication or presentation, please cite:

> **MultiQC: Summarize analysis results for multiple tools and samples in a single report**  
> *Philip Ewels, Måns Magnusson, Sverker Lundin and Max Käller*  
> Bioinformatics (2016)  
> doi: 10.1093/bioinformatics/btw354  
> PMID: 27312411

###### Save Settings

You can save the toolbox settings for this report to the browser.

 Save


---

###### Load Settings

Choose a saved report profile from the dropdown box below:

[ select ]

Load
 Delete
 Set default
 Clear default

###### About MultiQC

This report was generated using MultiQC, version 1.9

You can see a YouTube video describing how to use MultiQC reports here:
https://youtu.be/qPbIlO\_KWN0

For more information about MultiQC, including other videos and
extensive documentation, please visit http://multiqc.info

You can report bugs, suggest improvements and find the source code for MultiQC on GitHub:
https://github.com/ewels/MultiQC

MultiQC is published in Bioinformatics:

> **MultiQC: Summarize analysis results for multiple tools and samples in a single report**  
> *Philip Ewels, Måns Magnusson, Sverker Lundin and Max Käller*  
> Bioinformatics (2016)  
> doi: 10.1093/bioinformatics/btw354  
> PMID: 27312411

A modular tool to aggregate results from bioinformatics analyses across many samples into a single report.

###### JavaScript Disabled

MultiQC reports use JavaScript for plots and toolbox functions. It looks like
you have JavaScript disabled in your web browser. Please note that many of the report
functions will not work as intended.

Loading report..

Report
generated on 2021-05-13, 21:14
based on data in:
`/gpfs/home/fxr08zru/NER0008751/NER0008751_obj3_exp3_dmel/02_outputs/01_ovary/00_fastqc_raw_reads`

---

×
don't show again

**Welcome!** Not sure where to start?  
Watch a tutorial video
  *(6:06)*

#### General Statistics

 Copy table

 Configure Columns

 Sort by highlight

 Plot
Showing ^28^/_28_ rows and ^3^/_5_ columns.

| Sample Name | % Dups | % GC | Length | % Failed | M Seqs |
| --- | --- | --- | --- | --- | --- |
| 01\_NER0008751\_obj3\_exp3\_DC21\_ovary\_M\_RNA1\_rep1\_R1 | 77.5% | 51% | 100 bp | 27% | 56.4 |
| 01\_NER0008751\_obj3\_exp3\_DC21\_ovary\_M\_RNA1\_rep1\_R2 | 74.1% | 51% | 100 bp | 18% | 56.4 |
| 02\_NER0008751\_obj3\_exp3\_DC24\_ovary\_M\_RNA2\_rep1\_R1 | 72.1% | 52% | 100 bp | 27% | 31.8 |
| 02\_NER0008751\_obj3\_exp3\_DC24\_ovary\_M\_RNA2\_rep1\_R2 | 70.2% | 52% | 100 bp | 18% | 31.8 |
| 03\_NER0008751\_obj3\_exp3\_DC27\_ovary\_H\_RNA1\_rep1\_R1 | 77.2% | 52% | 100 bp | 27% | 57.3 |
| 03\_NER0008751\_obj3\_exp3\_DC27\_ovary\_H\_RNA1\_rep1\_R2 | 73.6% | 51% | 100 bp | 18% | 57.3 |
| 04\_NER0008751\_obj3\_exp3\_DC30\_ovary\_H\_RNA2\_rep1\_R1 | 75.0% | 52% | 100 bp | 27% | 46.5 |
| 04\_NER0008751\_obj3\_exp3\_DC30\_ovary\_H\_RNA2\_rep1\_R2 | 72.7% | 52% | 100 bp | 18% | 46.5 |
| 05\_NER0008751\_obj3\_exp3\_DC33\_ovary\_M\_RNA1\_rep2\_R1 | 78.2% | 52% | 100 bp | 27% | 67.3 |
| 05\_NER0008751\_obj3\_exp3\_DC33\_ovary\_M\_RNA1\_rep2\_R2 | 74.9% | 52% | 100 bp | 18% | 67.3 |
| 06\_NER0008751\_obj3\_exp3\_DC36\_ovary\_M\_RNA2\_rep2\_R1 | 73.6% | 51% | 100 bp | 27% | 41.8 |
| 06\_NER0008751\_obj3\_exp3\_DC36\_ovary\_M\_RNA2\_rep2\_R2 | 70.6% | 51% | 100 bp | 18% | 41.8 |
| 07\_NER0008751\_obj3\_exp3\_DC39\_ovary\_H\_RNA1\_rep2\_R1 | 74.7% | 52% | 100 bp | 27% | 40.9 |
| 07\_NER0008751\_obj3\_exp3\_DC39\_ovary\_H\_RNA1\_rep2\_R2 | 70.8% | 52% | 100 bp | 18% | 40.9 |
| 08\_NER0008751\_obj3\_exp3\_DC42\_ovary\_H\_RNA2\_rep2\_R1 | 77.9% | 51% | 100 bp | 27% | 63.6 |
| 08\_NER0008751\_obj3\_exp3\_DC42\_ovary\_H\_RNA2\_rep2\_R2 | 74.1% | 51% | 100 bp | 18% | 63.6 |
| 09\_NER0008751\_obj3\_exp3\_DC45\_ovary\_M\_RNA1\_rep3\_R1 | 77.9% | 52% | 100 bp | 27% | 67.4 |
| 09\_NER0008751\_obj3\_exp3\_DC45\_ovary\_M\_RNA1\_rep3\_R2 | 73.8% | 52% | 100 bp | 18% | 67.4 |
| 10\_NER0008751\_obj3\_exp3\_DC48\_ovary\_M\_RNA2\_rep3\_R1 | 72.5% | 52% | 100 bp | 27% | 35.2 |
| 10\_NER0008751\_obj3\_exp3\_DC48\_ovary\_M\_RNA2\_rep3\_R2 | 69.3% | 52% | 100 bp | 18% | 35.2 |
| 11\_NER0008751\_obj3\_exp3\_DC54\_ovary\_H\_RNA2\_rep3\_R1 | 76.3% | 52% | 100 bp | 27% | 55.2 |
| 11\_NER0008751\_obj3\_exp3\_DC54\_ovary\_H\_RNA2\_rep3\_R2 | 71.2% | 52% | 100 bp | 18% | 55.2 |
| 12\_NER0008751\_obj3\_exp3\_DC57\_ovary\_H\_RNA1\_rep4\_R1 | 77.8% | 52% | 100 bp | 27% | 65.4 |
| 12\_NER0008751\_obj3\_exp3\_DC57\_ovary\_H\_RNA1\_rep4\_R2 | 74.2% | 52% | 100 bp | 18% | 65.4 |
| 37\_NER0008751\_obj3\_exp3\_DC24\_ovary\_M\_RNA2\_rep1\_R1 | 87.8% | 52% | 100 bp | 27% | 218.3 |
| 37\_NER0008751\_obj3\_exp3\_DC24\_ovary\_M\_RNA2\_rep1\_R2 | 88.5% | 52% | 100 bp | 18% | 218.3 |
| 38\_NER0008751\_obj3\_exp3\_DC48\_ovary\_M\_RNA2\_rep3\_R1 | 88.2% | 52% | 100 bp | 27% | 243.6 |
| 38\_NER0008751\_obj3\_exp3\_DC48\_ovary\_M\_RNA2\_rep3\_R2 | 87.4% | 52% | 100 bp | 18% | 243.6 |

×

###### General Statistics: Columns

Uncheck the tick box to hide columns. Click and drag the handle on the left to change order.

Show All
Show None

| Sort | Visible | Group | Column | Description | ID | Scale |
| --- | --- | --- | --- | --- | --- | --- |
| \|\| |  | FastQC | % Dups | % Duplicate Reads | `percent_duplicates` | None |
| \|\| |  | FastQC | % GC | Average % GC Content | `percent_gc` | None |
| \|\| |  | FastQC | Length | Average Sequence Length (bp) | `avg_sequence_length` | None |
| \|\| |  | FastQC | % Failed | Percentage of modules failed in FastQC report (includes those not plotted here) | `percent_fails` | None |
| \|\| |  | FastQC | M Seqs | Total Sequences (millions) | `total_sequences` | read\_count |

Close

#### FastQC

FastQC is a quality control tool for high throughput sequence data, written by Simon Andrews at the Babraham Institute in Cambridge.

##### Sequence Counts Help

Sequence counts for each sample. Duplicate read counts are an estimate only.

This plot show the total number of reads, broken down into unique and duplicate
if possible (only more recent versions of FastQC give duplicate info).

You can read more about duplicate calculation in the
FastQC documentation.
A small part has been copied here for convenience:

*Only sequences which first appear in the first 100,000 sequences
in each file are analysed. This should be enough to get a good impression
for the duplication levels in the whole file. Each sequence is tracked to
the end of the file to give a representative count of the overall duplication level.*

*The duplication detection requires an exact sequence match over the whole length of
the sequence. Any reads over 75bp in length are truncated to 50bp for this analysis.*

Number of reads
Percentages

loading..

---

##### Sequence Quality Histograms Help

The mean quality value across each base position in the read.

To enable multiple samples to be plotted on the same graph, only the mean quality
scores are plotted (unlike the box plots seen in FastQC reports).

Taken from the FastQC help:

*The y-axis on the graph shows the quality scores. The higher the score, the better
the base call. The background of the graph divides the y axis into very good quality
calls (green), calls of reasonable quality (orange), and calls of poor quality (red).
The quality of calls on most platforms will degrade as the run progresses, so it is
common to see base calls falling into the orange area towards the end of a read.*

loading..

---

##### Per Sequence Quality Scores Help

The number of reads with average quality scores. Shows if a subset of reads has poor quality.

From the FastQC help:

*The per sequence quality score report allows you to see if a subset of your
sequences have universally low quality values. It is often the case that a
subset of sequences will have universally poor quality, however these should
represent only a small percentage of the total sequences.*

loading..

---

##### Per Base Sequence Content Help

The proportion of each base position for which each of the four normal DNA bases has been called.

To enable multiple samples to be shown in a single plot, the base composition data
is shown as a heatmap. The colours represent the balance between the four bases:
an even distribution should give an even muddy brown colour. Hover over the plot
to see the percentage of the four bases under the cursor.

**To see the data as a line plot, as in the original FastQC graph, click on a sample track.**

From the FastQC help:

*Per Base Sequence Content plots out the proportion of each base position in a
file for which each of the four normal DNA bases has been called.*

*In a random library you would expect that there would be little to no difference
between the different bases of a sequence run, so the lines in this plot should
run parallel with each other. The relative amount of each base should reflect
the overall amount of these bases in your genome, but in any case they should
not be hugely imbalanced from each other.*

*It's worth noting that some types of library will always produce biased sequence
composition, normally at the start of the read. Libraries produced by priming
using random hexamers (including nearly all RNA-Seq libraries) and those which
were fragmented using transposases inherit an intrinsic bias in the positions
at which reads start. This bias does not concern an absolute sequence, but instead
provides enrichement of a number of different K-mers at the 5' end of the reads.
Whilst this is a true technical bias, it isn't something which can be corrected
by trimming and in most cases doesn't seem to adversely affect the downstream
analysis.*

Click a sample row to see a line plot for that dataset.

###### Rollover for sample name

 Export Plot

Position: -

%T: -

%C: -

%A: -

%G: -

---

##### Per Sequence GC Content Help

The average GC content of reads. Normal random library typically have a
roughly normal distribution of GC content.

From the FastQC help:

*This module measures the GC content across the whole length of each sequence
in a file and compares it to a modelled normal distribution of GC content.*

*In a normal random library you would expect to see a roughly normal distribution
of GC content where the central peak corresponds to the overall GC content of
the underlying genome. Since we don't know the the GC content of the genome the
modal GC content is calculated from the observed data and used to build a
reference distribution.*

*An unusually shaped distribution could indicate a contaminated library or
some other kinds of biased subset. A normal distribution which is shifted
indicates some systematic bias which is independent of base position. If there
is a systematic bias which creates a shifted normal distribution then this won't
be flagged as an error by the module since it doesn't know what your genome's
GC content should be.*

Percentages
Counts

loading..

---

##### Per Base N Content Help

The percentage of base calls at each position for which an `N` was called.

From the FastQC help:

*If a sequencer is unable to make a base call with sufficient confidence then it will
normally substitute an `N` rather than a conventional base call. This graph shows the
percentage of base calls at each position for which an `N` was called.*

*It's not unusual to see a very low proportion of Ns appearing in a sequence, especially
nearer the end of a sequence. However, if this proportion rises above a few percent
it suggests that the analysis pipeline was unable to interpret the data well enough to
make valid base calls.*

loading..

---

##### Sequence Length Distribution

All samples have sequences of a single length (100bp).

---

##### Sequence Duplication Levels Help

The relative level of duplication found for every sequence.

From the FastQC Help:

*In a diverse library most sequences will occur only once in the final set.
A low level of duplication may indicate a very high level of coverage of the
target sequence, but a high level of duplication is more likely to indicate
some kind of enrichment bias (eg PCR over amplification). This graph shows
the degree of duplication for every sequence in a library: the relative
number of sequences with different degrees of duplication.*

*Only sequences which first appear in the first 100,000 sequences
in each file are analysed. This should be enough to get a good impression
for the duplication levels in the whole file. Each sequence is tracked to
the end of the file to give a representative count of the overall duplication level.*

*The duplication detection requires an exact sequence match over the whole length of
the sequence. Any reads over 75bp in length are truncated to 50bp for this analysis.*

*In a properly diverse library most sequences should fall into the far left of the
plot in both the red and blue lines. A general level of enrichment, indicating broad
oversequencing in the library will tend to flatten the lines, lowering the low end
and generally raising other categories. More specific enrichments of subsets, or
the presence of low complexity contaminants will tend to produce spikes towards the
right of the plot.*

loading..

---

##### Overrepresented sequences Help

The total amount of overrepresented sequences found in each library.

FastQC calculates and lists overrepresented sequences in FastQ files. It would not be
possible to show this for all samples in a MultiQC report, so instead this plot shows
the *number of sequences* categorized as over represented.

Sometimes, a single sequence may account for a large number of reads in a dataset.
To show this, the bars are split into two: the first shows the overrepresented reads
that come from the single most common sequence. The second shows the total count
from all remaining overrepresented sequences.

From the FastQC Help:

*A normal high-throughput library will contain a diverse set of sequences, with no
individual sequence making up a tiny fraction of the whole. Finding that a single
sequence is very overrepresented in the set either means that it is highly biologically
significant, or indicates that the library is contaminated, or not as diverse as you expected.*

*FastQC lists all of the sequences which make up more than 0.1% of the total.
To conserve memory only sequences which appear in the first 100,000 sequences are tracked
to the end of the file. It is therefore possible that a sequence which is overrepresented
but doesn't appear at the start of the file for some reason could be missed by this module.*

28 samples had less than 1% of reads made up of overrepresented sequences

---

##### Adapter Content Help

The cumulative percentage count of the proportion of your
library which has seen each of the adapter sequences at each position.

Note that only samples with ≥ 0.1% adapter contamination are shown.

There may be several lines per sample, as one is shown for each adapter
detected in the file.

From the FastQC Help:

*The plot shows a cumulative percentage count of the proportion
of your library which has seen each of the adapter sequences at each position.
Once a sequence has been seen in a read it is counted as being present
right through to the end of the read so the percentages you see will only
increase as the read length goes on.*

loading..

---

##### Status Checks Help

Status for each FastQC section showing whether results seem entirely normal (green),
slightly abnormal (orange) or very unusual (red).

FastQC assigns a status for each section of the report.
These give a quick evaluation of whether the results of the analysis seem
entirely normal (green), slightly abnormal (orange) or very unusual (red).

It is important to stress that although the analysis results appear to give a pass/fail result,
these evaluations must be taken in the context of what you expect from your library.
A 'normal' sample as far as FastQC is concerned is random and diverse.
Some experiments may be expected to produce libraries which are biased in particular ways.
You should treat the summary evaluations therefore as pointers to where you should concentrate
your attention and understand why your library may not look random and diverse.

Specific guidance on how to interpret the output of each module can be found in the relevant
report section, or in the FastQC help.

In this heatmap, we summarise all of these into a single heatmap for a quick overview.
Note that not all FastQC sections have plots in MultiQC reports, but all status checks
are shown in this heatmap.

Sort by highlight

loading..

**MultiQC v1.9** 
- Written by Phil Ewels,
available on GitHub.

This report uses HighCharts,
jQuery,
jQuery UI,
Bootstrap,
FileSaver.js and
clipboard.js.

×

##### Plot Table Data

Select Column

Select Column

Please select two table columns.

Close

×

##### Regex Help

Toolbox search strings can behave as regular expressions (regexes). Click a button below to see an example of it in action. Try modifying them yourself in the text box.

`^` (start of string)
`$` (end of string)
`[]` (character choice)
`\d` (shorthand for `[0-9]`)
`\w` (shorthand for `[0-9a-zA-Z_]`)
`.` (any character)
`\.` (literal full stop)
`()` `|` (group / separator)
`*` (prev char 0 or more)
`+` (prev char 1 or more)
`?` (prev char 0 or 1)
`{}` (char num times)
`{,}` (count range)

```
samp_1
samp_1_edited
samp_2
samp_2_edited
samp_3
samp_3_edited
prepended_samp_1
tmp_samp_1_edited
tmpp_samp_1_edited
tmppp_samp_1_edited
#samp_1_edited.tmp
samp_11
samp_11111
```

See regex101.com for a more heavy duty testing suite.

Close
