## Supplemental file S1 for "Developmental diet alters the fecundity-longevity relationship and age-related gene expression in *Drosophila melanogaster*"

MultiQC Report


### Toggle navigation v1.9

Loading report..

- General Stats
- FastQC
  - Sequence Counts
  - Sequence Quality Histograms
  - Per Sequence Quality Scores
  - Per Base Sequence Content
  - Per Sequence GC Content
  - Per Base N Content
  - Sequence Length Distribution
  - Sequence Duplication Levels
  - Overrepresented sequences
  - Adapter Content
  - Status Checks

Toolbox

##### MultiQC Toolbox

###### Apply Highlight Samples

+

Regex mode off
help
 Clear

###### Apply Rename Samples

+

Click here for bulk input.

Paste two columns of a tab-delimited table here (eg. from Excel).

First column should be the old name, second column the new name.

Add

Regex mode off
help
 Clear

###### Apply Show / Hide Samples

Hide matching samples

Show only matching samples

+

Regex mode off
help
 Clear

###### Export Plots

- Images
- Data

px

px

Aspect ratio

PNG
JPEG
SVG

Plot scaling

X

Download the raw data used to create the plots in this report below:

Format:

Tab-separated
Comma-separated
JSON

Note that additional data was saved in `00_NER0008751_obj3_exp3_supplementary_file_S1_head_fastqc_multiqc_data` when this report was generated.

---

###### Choose Plots

 All
 None


---


   Download Plot Images

If you use plots from MultiQC in a publication or presentation, please cite:

> **MultiQC: Summarize analysis results for multiple tools and samples in a single report**  
> *Philip Ewels, Måns Magnusson, Sverker Lundin and Max Käller*  
> Bioinformatics (2016)  
> doi: 10.1093/bioinformatics/btw354  
> PMID: 27312411

###### Save Settings

You can save the toolbox settings for this report to the browser.

 Save


---

###### Load Settings

Choose a saved report profile from the dropdown box below:

[ select ]

Load
 Delete
 Set default
 Clear default

###### About MultiQC

This report was generated using MultiQC, version 1.9

You can see a YouTube video describing how to use MultiQC reports here:
https://youtu.be/qPbIlO\_KWN0

For more information about MultiQC, including other videos and
extensive documentation, please visit http://multiqc.info

You can report bugs, suggest improvements and find the source code for MultiQC on GitHub:
https://github.com/ewels/MultiQC

MultiQC is published in Bioinformatics:

> **MultiQC: Summarize analysis results for multiple tools and samples in a single report**  
> *Philip Ewels, Måns Magnusson, Sverker Lundin and Max Käller*  
> Bioinformatics (2016)  
> doi: 10.1093/bioinformatics/btw354  
> PMID: 27312411

A modular tool to aggregate results from bioinformatics analyses across many samples into a single report.

###### JavaScript Disabled

MultiQC reports use JavaScript for plots and toolbox functions. It looks like
you have JavaScript disabled in your web browser. Please note that many of the report
functions will not work as intended.

Loading report..

Report
generated on 2021-05-13, 20:14
based on data in:
`/gpfs/home/fxr08zru/NER0008751/NER0008751_obj3_exp3_dmel/02_outputs/02_head/00_fastqc_raw_reads`

---

×
don't show again

**Welcome!** Not sure where to start?  
Watch a tutorial video
  *(6:06)*

#### General Statistics

 Copy table

 Configure Columns

 Sort by highlight

 Plot
Showing ^24^/_24_ rows and ^3^/_5_ columns.

| Sample Name | % Dups | % GC | Length | % Failed | M Seqs |
| --- | --- | --- | --- | --- | --- |
| 13\_NER0008751\_obj3\_exp3\_DC19\_head\_M\_RNA1\_rep1\_R1 | 80.7% | 51% | 100 bp | 27% | 61.0 |
| 13\_NER0008751\_obj3\_exp3\_DC19\_head\_M\_RNA1\_rep1\_R2 | 74.3% | 51% | 100 bp | 18% | 61.0 |
| 14\_NER0008751\_obj3\_exp3\_DC22\_head\_M\_RNA2\_rep1\_R1 | 74.6% | 50% | 100 bp | 27% | 47.9 |
| 14\_NER0008751\_obj3\_exp3\_DC22\_head\_M\_RNA2\_rep1\_R2 | 69.9% | 49% | 100 bp | 18% | 47.9 |
| 15\_NER0008751\_obj3\_exp3\_DC25\_head\_H\_RNA1\_rep1\_R1 | 79.3% | 51% | 100 bp | 27% | 68.0 |
| 15\_NER0008751\_obj3\_exp3\_DC25\_head\_H\_RNA1\_rep1\_R2 | 74.4% | 51% | 100 bp | 18% | 68.0 |
| 16\_NER0008751\_obj3\_exp3\_DC28\_head\_H\_RNA2\_rep1\_R1 | 77.5% | 50% | 100 bp | 27% | 78.7 |
| 16\_NER0008751\_obj3\_exp3\_DC28\_head\_H\_RNA2\_rep1\_R2 | 73.9% | 50% | 100 bp | 18% | 78.7 |
| 17\_NER0008751\_obj3\_exp3\_DC31\_head\_M\_RNA1\_rep2\_R1 | 78.6% | 51% | 100 bp | 27% | 70.7 |
| 17\_NER0008751\_obj3\_exp3\_DC31\_head\_M\_RNA1\_rep2\_R2 | 74.7% | 50% | 100 bp | 18% | 70.7 |
| 18\_NER0008751\_obj3\_exp3\_DC34\_head\_M\_RNA2\_rep2\_R1 | 74.1% | 49% | 100 bp | 27% | 46.7 |
| 18\_NER0008751\_obj3\_exp3\_DC34\_head\_M\_RNA2\_rep2\_R2 | 71.1% | 49% | 100 bp | 27% | 46.7 |
| 19\_NER0008751\_obj3\_exp3\_DC37\_head\_H\_RNA1\_rep2\_R1 | 75.5% | 51% | 100 bp | 27% | 46.9 |
| 19\_NER0008751\_obj3\_exp3\_DC37\_head\_H\_RNA1\_rep2\_R2 | 70.3% | 50% | 100 bp | 18% | 46.9 |
| 20\_NER0008751\_obj3\_exp3\_DC43\_head\_M\_RNA1\_rep3\_R1 | 81.0% | 51% | 100 bp | 27% | 58.3 |
| 20\_NER0008751\_obj3\_exp3\_DC43\_head\_M\_RNA1\_rep3\_R2 | 77.3% | 51% | 100 bp | 18% | 58.3 |
| 21\_NER0008751\_obj3\_exp3\_DC46\_head\_M\_RNA2\_rep3\_R1 | 77.4% | 49% | 100 bp | 27% | 80.2 |
| 21\_NER0008751\_obj3\_exp3\_DC46\_head\_M\_RNA2\_rep3\_R2 | 71.8% | 49% | 100 bp | 18% | 80.2 |
| 22\_NER0008751\_obj3\_exp3\_DC49\_head\_H\_RNA1\_rep3\_R1 | 81.3% | 50% | 100 bp | 27% | 117.6 |
| 22\_NER0008751\_obj3\_exp3\_DC49\_head\_H\_RNA1\_rep3\_R2 | 76.5% | 50% | 100 bp | 18% | 117.6 |
| 23\_NER0008751\_obj3\_exp3\_DC52\_head\_H\_RNA2\_rep3\_R1 | 77.9% | 50% | 100 bp | 27% | 84.8 |
| 23\_NER0008751\_obj3\_exp3\_DC52\_head\_H\_RNA2\_rep3\_R2 | 74.3% | 50% | 100 bp | 18% | 84.8 |
| 24\_NER0008751\_obj3\_exp3\_DC58\_head\_H\_RNA2\_rep4\_R1 | 79.6% | 50% | 100 bp | 27% | 97.7 |
| 24\_NER0008751\_obj3\_exp3\_DC58\_head\_H\_RNA2\_rep4\_R2 | 76.4% | 50% | 100 bp | 18% | 97.7 |

Close
